# HYL001, a new potent TGFβ signaling inhibitor that is efficacious against microsatellite stable CRC metastasis in combination with immune checkpoint therapy in mice

**DOI:** 10.1101/2024.05.10.593510

**Authors:** Daniele V. F. Tauriello, Elena Sancho, Daniel Byrom, Carolina Sanchez-Zarzalejo, Maria Salvany, Ana Henriques, Sergio Palomo-Ponce, Marta Sevillano, Xavier Hernando-Momblona, Joan A. Matarin, Israel Ramos, Irene Ruano, Neus Prats, Eduard Batlle, Antoni Riera

## Abstract

Blockade of the TGFβ signalling pathway has emerged from preclinical studies as a potential treatment to enhance the efficacy of immune checkpoint inhibition in advanced colorectal cancer (CRC) and several other types of cancer. However, clinical translation of first-generation inhibitors has known little success. Here, we report the synthesis and characterization of HYL001, a potent inhibitor of TGFβ receptor 1 (ALK5), that is approximately 9 times more efficacious than the structurally related compound galunisertib, while maintaining a favourable safety profile. HYL001 in combination with immune checkpoint blockade (anti-PD1) eradicates liver metastases generated in mice by microsatellite stable, aggressive colorectal cancer tumours at doses where galunisertib is ineffective.

**GRAPHICAL ABSTRACT:** 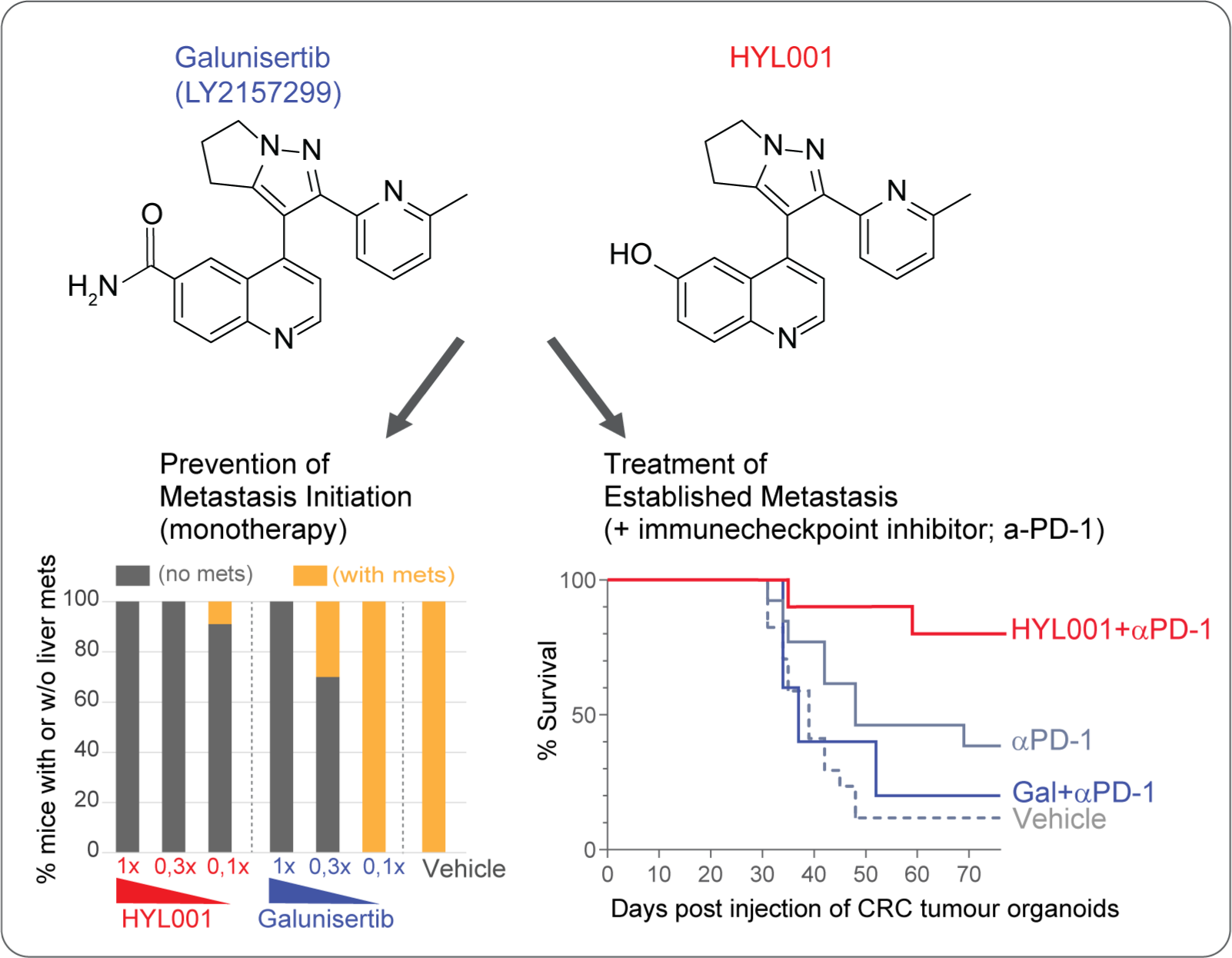

## INTRODUCTION

Cancer takes almost 10 million lives each year^1^. Advanced disease with tumours spreading to distant organs (i.e. metastasis) is associated with most cancer deaths. For example, despite considerable progress, metastatic colorectal cancer (mCRC), still has a low 5-year survival rate (<20%) and is thus in urgent need for better treatment options. Among promising recent developments in cancer treatment are immunotherapies, notably including immune checkpoint inhibitors (ICIs)^2,3^. However, many patients remain unresponsive to immune therapies. In the case of mCRC, responses seem limited to a small fraction of patients who have tumours that are classified as mismatch-repair deficient or microsatellite-instable (dMMR/MSI). These typically have a high tumour mutational burden that can elicit specific immune responses. Boosting the efficacy of immunotherapy to larger groups of patients would represent a clinical advance with a tremendous impact.

The past decades, fundamental evidence has mounted for TGFβ signalling as an immunosuppressive pathway with key relevance in cancer^4^. Indeed, tumours displaying a TGFβ-activated stroma represent a clinical entity associated with poor prognosis and immune evasion^5–9^. In 2018, we and others showed that increased activity of TGFβ in the tumour microenvironment (TME) is associated with lack of response to anti-PD-1/anti-PD-L1 in preclinical CRC models and in human metastatic urothelial cancer^5,10^. We demonstrated that TGFβ blockade synergizes with immune checkpoint blockade to cure mice with multiple colorectal cancer liver metastases^5^. Interestingly, our murine mCRC model is mismatch-repair proficient (pMMR), microsatellite stable (MSS) and represents the majority of tumours that— especially when KRAS-mutant—have not seen many new efficacious treatment options recently. Additional corroborating studies have been reported since, and clinical trials have begun testing this or similar TGFβ inhibition-based immuno-oncology strategies in a range of cancer types^6,7^. Furthermore, TGFβ blockade could also be useful to treat additional diseases where this cytokine plays a role, such as fibrosis in vital organs like lung, liver, kidney and skin^11,12^.

Despite all this, and the development of various agents to inhibit TGFβ signalling, there are no clinically approved drugs to inhibit this pathway^6,13^. The development of TGFβ inhibitors has been hampered in part by their associations with on-target cardiac toxicity in rats^14^, which may likely translate to humans and therefore poses limits for drug dosing. As an example, from **1**, a class of quinoline-substituted dihydropyrrolopyrazoles **1a** (galunisertib, LY2157299) is a small molecule inhibitor that targets the TGFβ signaling pathway by competitively binding the ATP pocket of the TGFβ receptor type I (TGFBR1; ALK5)^15,16^. Given its relatively low toxicity profile, galunisertib has been extensively tested in clinical oncology studies to inhibit TGFβ signaling^6,17^. This favourable safety profile appears related to a relatively low potency. Accordingly, the aforementioned efficacy against murine CRC liver metastasis required elevated dosing of this drug^5^.

An efficacious murine dose for **1a** was established in our model at 720–800 mg/kg twice daily (BID), which is markedly higher than previously reported^15^. Although our dosing came with practical challenges, it caused only minor adverse effects in mice. Yet, such elevated doses would very likely not be tolerable in humans. We therefore sought to design a new TGFβ inhibitor with a better potency–safety profile. Furthermore, we aimed for the improved molecule to have a ‘handle’ to which we could attach a variety of groups, enabling further tuning of pharmacochemical properties. Moreover, this could enable a caged prodrug with spatiotemporally controlled release. In parallel, other second-generation ALK5 kinase inhibitors have been developed that appear to maintain acceptable toxicity despite higher potency. One of these, **2** (vactosertib, TEW-7197)^18^, is currently undergoing clinical investigation for treatment of various solid tumours^6,13^.

## RESULTS

### Chemical synthesis of HYL001

In 2008, Li *et. al.* published a structure activity relationship study on the quinoline domain of the **1** series of ALK5 inhibitors^16^. From this study, it was apparent that the substitution on the 2-position had a negative impact on the IC_50_ value when changed for anything but a hydrogen atom. On the other hand, derivatisation at the 7-position produced some potent inhibitors. Interestingly, a hydroxyl group at 7-position (7-OH, R_2_ = H) afforded a clearly less potent compound (IC_50_, 160 nM) than galunisertib (51 nM)^17^. However, modification at the 6-position appeared to be a promising starting point for us, as the 6-bromo compound (6-Br, R_2_ = CH_3_) gave a lower IC_50_ (89 nM). Therefore, we synthesized this previously unreported compound with a hydroxyl group at the 6-position **1b**, which we called HYL001 (Figure 1).

**Figure 1.**
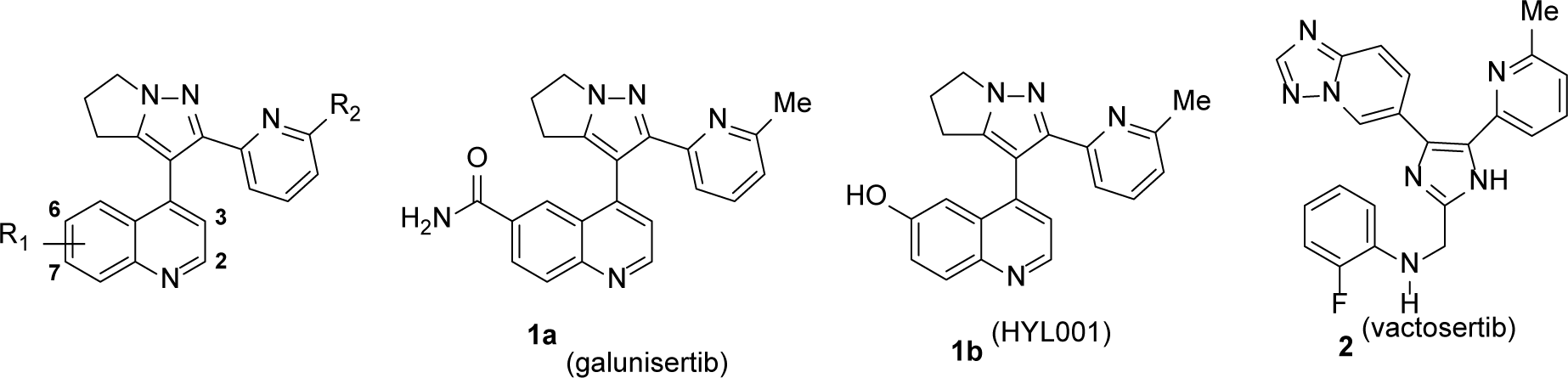
Chemical structures of ALK5 inhibitors, showing galunisertib (**1a**), HYL001 (**1b**) and vactosertib (**2**).

The synthesis of HYL001 was based on the same approach as galunisertib. The target compound should be prepared from compounds bearing a functional group X on the 6-position that could be transformed into the desired phenol. These compounds would be prepared from the corresponding 6-substituted-4-methylquinolines (**3**), easily accessible by a Doebner-Miller reaction of methyl vinyl ketone and the corresponding aniline (Figure 2).

**Figure 2.**
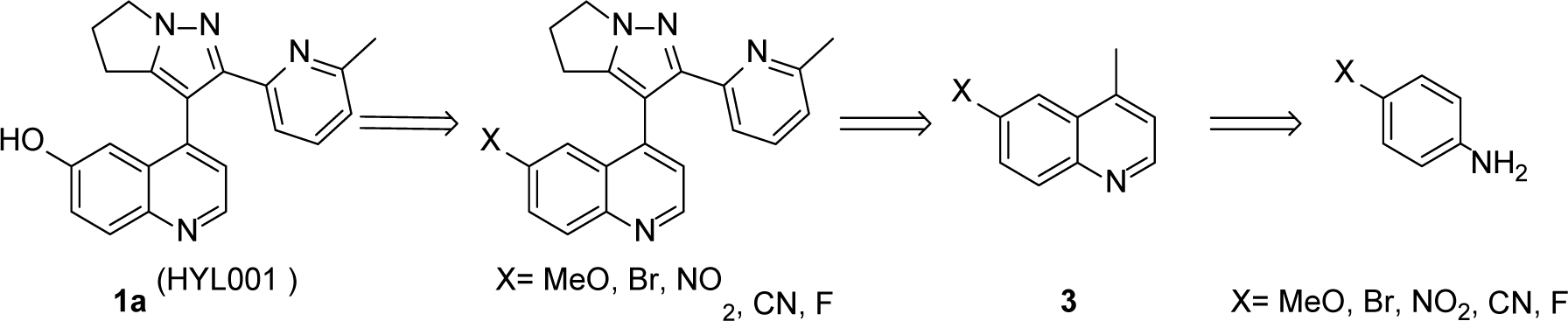
Retrosynthetic analysis of HYL001.

Although the use of nitro, fluoro and cyanoquinolines was explored, none of these approaches gave good yields of the desired product. Gratifyingly, the use of a methoxy group as a protected phenol was successful, allowing us the preparation of HYL001 (Scheme 1). However, the 6-methoxy-4-methylquinoline (**3a**) was an inconvenient starting material. The yield of the Doebner-Miller reaction was low (41%), the product was difficult to purify due to a large amount of ferric salts (chromatography was needed), and LDA was necessary for the alkylation with methyl 6-methylpicolinate to give compound **4a**. Nevertheless, the construction of the dihydropyrrolopyrazole ring to give **5a** as well as the deprotection of the methyl ether went as expected affording **1a** (HYL001) in good overall yield (Scheme 1).

The synthesis of HYL001 was next approached starting from 4-bromoaniline (Scheme 2). Its preparation by the Doebner-Miller reaction was more convenient since it could be done without the need of FeCl_3_, and the purification greatly improved. The subsequent alkylation was performed using HMDS as base, affording **3b** in excellent yield. Preparation of **4b** took place uneventfully. However, the conversion of the bromo derivative **4b** into the phenol was more difficult than expected. We eventually found that **5b** could be converted into boronate **6** by a Miyaura borilation. Oxidation and hydrolysis of the boronate gave the desired HYL001 in good yield. However, the procedure required a chromatographic purification of **5b** that prevented large scale synthesis. The procedure could be further improved by the direct conversion of **5b** into the final compound using tris(dibenzylideneacetone)dipalladium(0) and tetramethyl di-tBuXPhos as ligand. In this way a convenient and scalable procedure allowed the preparation of multigram quantities of HYL001 without any chromatographic purification.

**Scheme 1.**
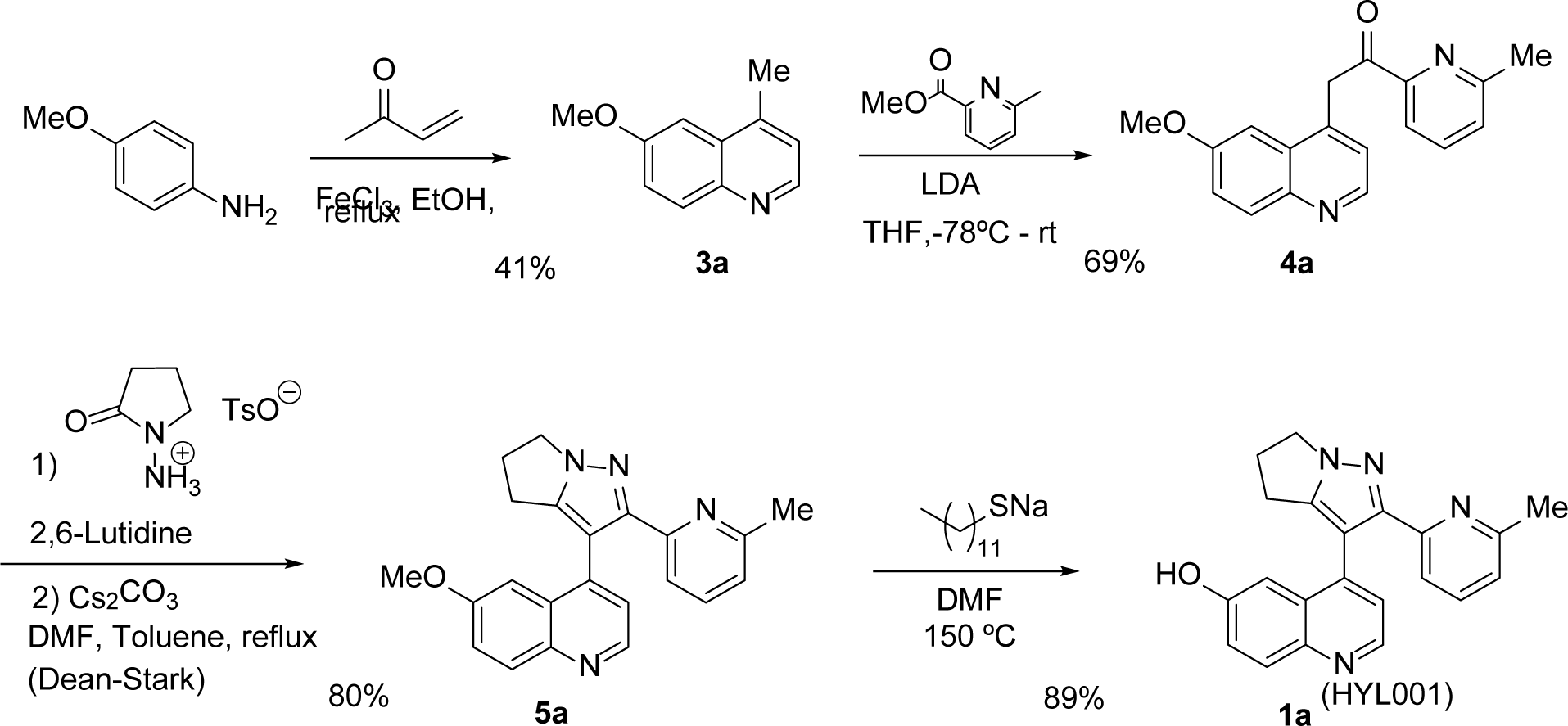
Synthesis of HYL001 from 4-methoxyaniline.

**Scheme 2.**
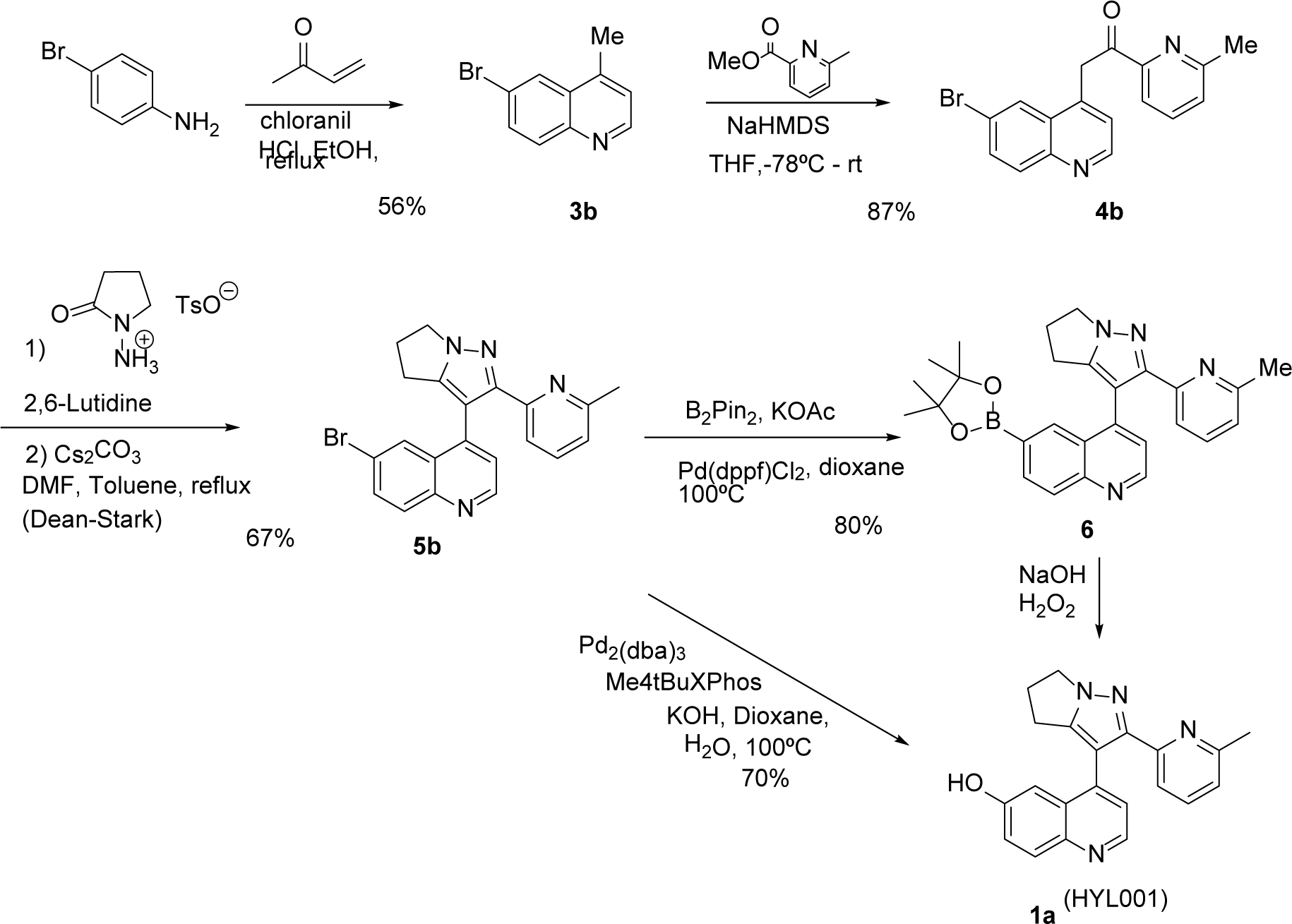
Synthesis of HYL001 from 4-bromoaniline.

### Chemical synthesis of HYL002

As a proof-of-concept prodrug that allows for soluble formulation, facilitating administration at least in the preclinical setting, we leveraged the hydroxyl handle to design HYL002. We added a carbolnyl-4-piperidinopiperidine group, in analogy to the ester that solubilizes SN-38 into irinotecan, and that can be cleaved via hydrolysis inside cells^19^ (Figure 3).

**Figure 3.**
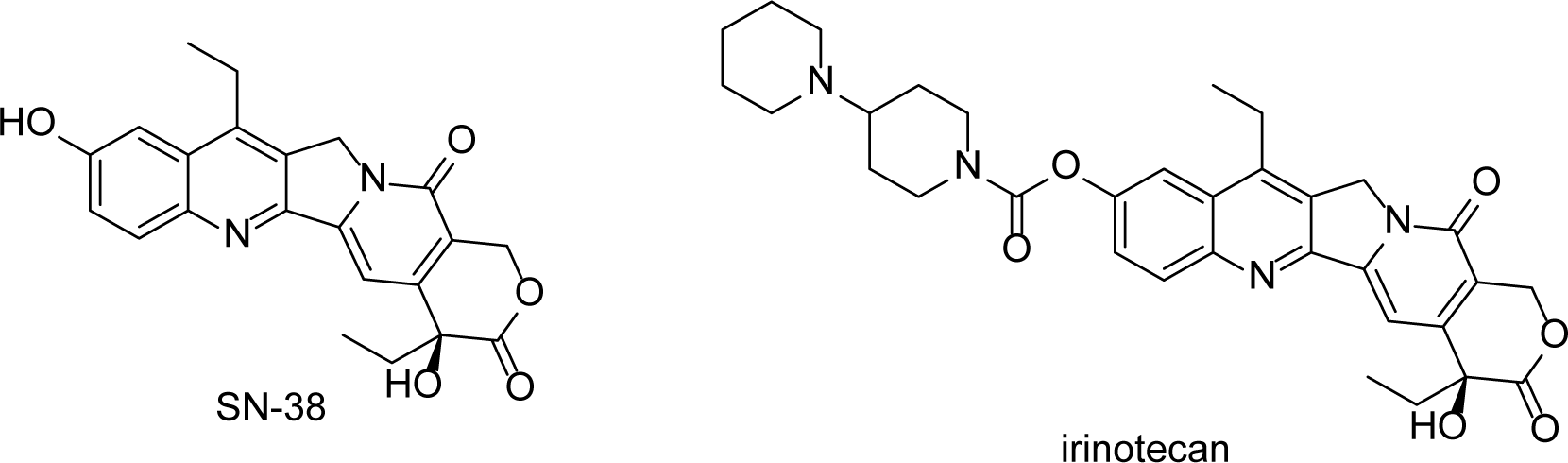
Irinotecan as the carbonyl-4-piperidinopiperidine ester of SN-38.

HYL001 reacted with commercially available 4-piperidinopiperidine-1-carbonyl chloride in chloroform using triethylamine and catalytic amounts of DMAP (Scheme 3). We then transformed the resulting compound HYL002 into the corresponding HCl salt to aid *in vivo* drug formulation. The free base was dissolved in dioxane under N_2_, where it was treated dropwise with 10 equivalents of 4M HCl in dioxane at room temperature with vigorous stirring. The reaction mixture was then taken to dryness *in vacuo* giving the HCl salt of HYL002.

**Scheme 3.**
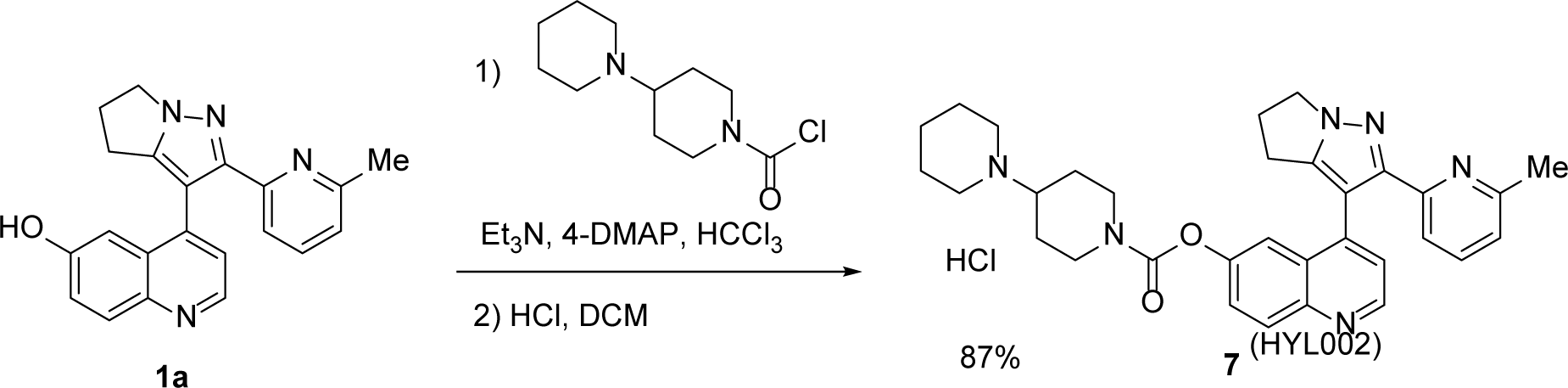
Synthesis of pro-drug **HYL002**

### Selectivity profile of HYL001

The *in vitro* binding affinity of HYL001 towards selected members of the TGFβ receptor family was assessed using a competitive binding assay. HYL001 is a potent ALK4 and ALK5 inhibitor with a K_d_ of 13 nM and 22.5 nM, respectively (Table 1). In comparison, the reported values for galunisertib are around 78 nM and 172 nM, respectively^16^. However, in the same assay as performed for HYL001, we found a K_d_ of 52 nM for galunisertib against ALK5. We also included vactosertib, which gave an ALK5 K_d_ around 4 nM (Table 1).

**Table 1.**
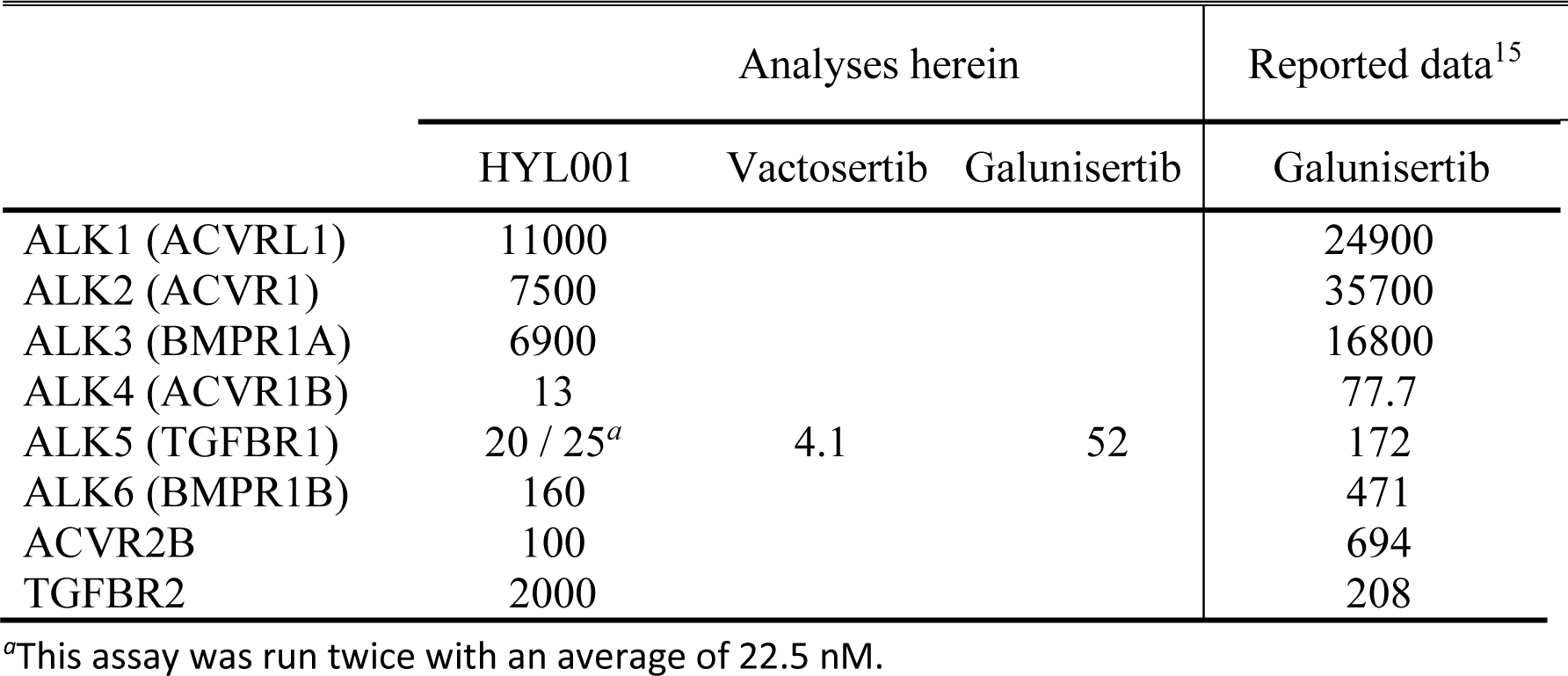
HYL001 selectivity towards ALK family members. Binding constants (K_d_) in nM.

|  | Analyses herein |  |  | Reported data<sup>15</sup> |
| --- | --- | --- | --- | --- |
|  | HYL001 | Vactosertib | Galunisertib | Galunisertib |
| ALK1 (ACVRL1) | 11000 |  |  | 24900 |
| ALK2 (ACVR1) | 7500 |  |  | 35700 |
| ALK3 (BMPR1A) | 6900 |  |  | 16800 |
| ALK4 (ACVR1B) | 13 |  |  | 77.7 |
| ALK5 (TGFB1) | 20 / 25<sup>a</sup> | 4.1 | 52 | 172 |
| ALK6 (BMPR1B) | 160 |  |  | 471 |
| ACVR2B | 100 |  |  | 694 |
| TGFB2 | 2000 |  |  | 208 |

<sup>a</sup>This assay was run twice with an average of 22.5 nM.

We also assessed selectivity of these three inhibitors towards a panel of 97 wild-type and disease-relevant mutant kinases, distributed throughout various kinase families. An initial analysis at high concentration indicated possible interactions with 13 of 90 kinases tested (>65% competitive binding inhibition by HYL001 at 10 μM) and with mutants of 2 additional kinases (Figure 4A). However, at a lower concentration (300 nM), the only interaction left was with p38α.

**Figure 4.**
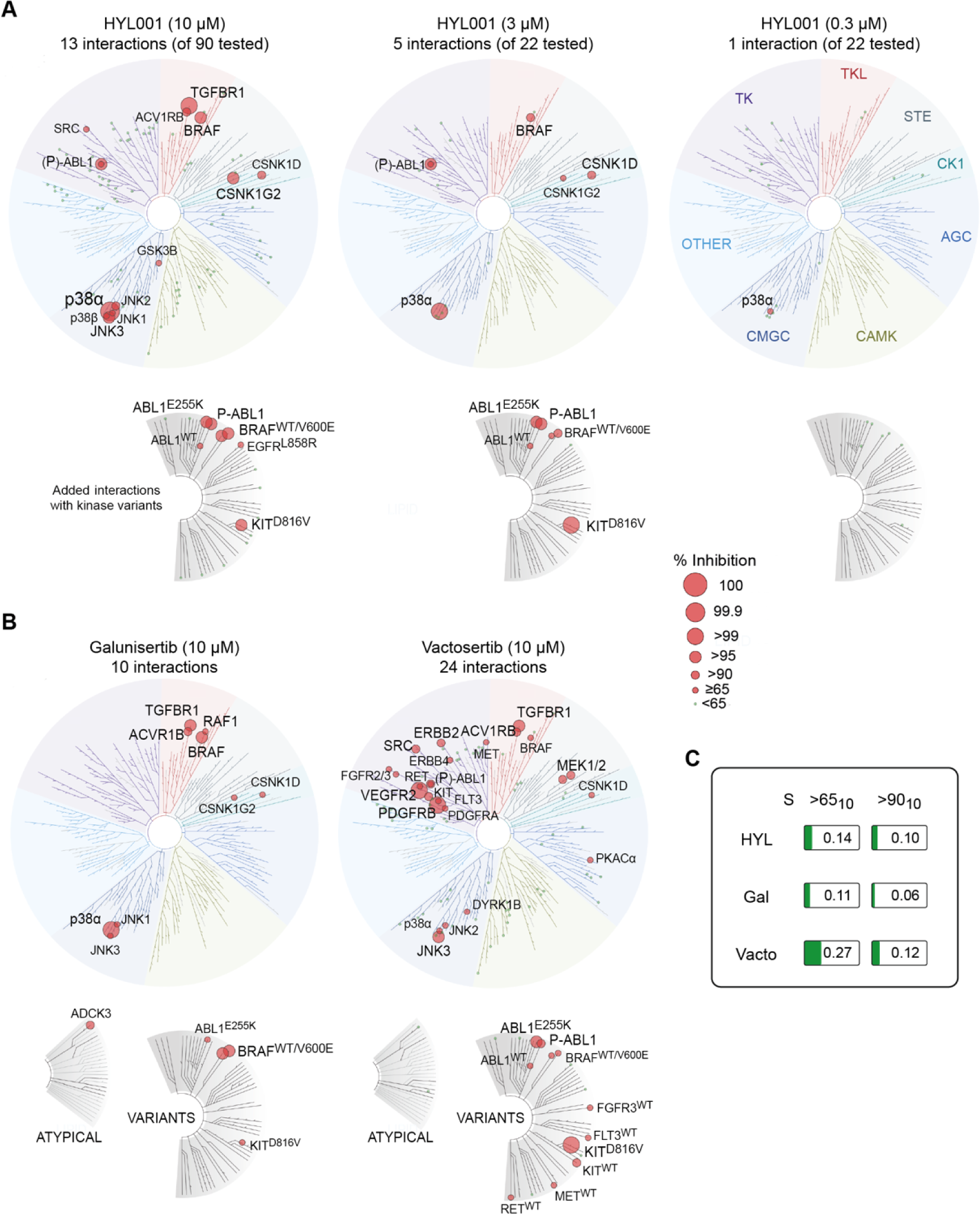
Selectivity of HYL001. **(A)** Data for HYL001 towards a panel of 90 kinases (and 7 kinase variants) at 10 μM. 22 Kinases inhibiting >50% (excluding those from Table 1) were subsequently tested at 3 μM and 300 nM. Visualized using TREE*spot*: interactions are depicted by red dots, with size proportional to inhibition of control ligand binding (shown if >65%). (**B**) Data for galunisertib and vactosertib at 10 μM towards the larger panel (N = 97). (**C**) Selectivity scores (fraction of wildtype kinases inhibited >65% or >90% at 10 μM compound concentration). *TK: tyrosine kinases; TKL: tyrosine kinase-like kinases; STE: serine/threonine kinases; CK1: casein kinase 1 family; AGC: serine/threonine kinases, regulated by secondary messengers such as cyclic AMP (PKA) or lipids (PKC); CAMK: Ca^2+^/calmodulin-dependent protein kinase family; CMGC: primarily proline-directed serine/threonine kinases (CDKs, GSKs, MAP kinases and CDK-like kinases)*.

Calculting the fraction of non-mutant target kinases with >65% inhibition at 10 μM, S(>65_10_) HYL001 scored 0.144. Its selectivity score for >90% inhibition S(>90_10_) was 0.1, which indicates a slightly lower selectivity compared to galunisertib, but higher selectivity than vactosertib (Figure 4B, C).

### *In vitro* enzymatic activity

ALK5 IC_50_ values for HYL001, galunisertib and vactosertib were obtained in a kinase activity assay with recombinant proteins (Table 2). Compared to galunisertib, the value for HYL001 of 177 nM constitutes an approximate 2.5-fold increase—in accordance with the increase in binding affinity (Table 1). This IC_50_ of HYL001 is around 3.5-fold higher than that of vactosertib in this assay (Table 2). We also analyzed the kinase inhibition activity of HYL001 towards p38α (Table 2). HYL001 is stronger inhibitor of p38α than galunisertib. Vactosertib, on the contrary, shows little activity towards this kinase as it was specifically selected to not inhibit p38α^18^.

**Table 2.**
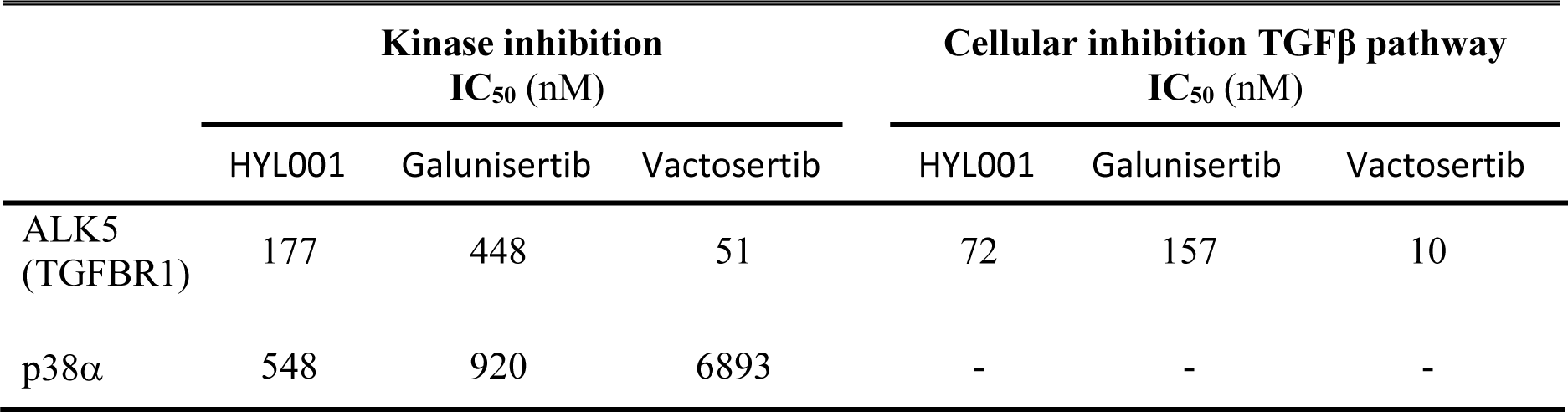
In vitro kinase inhibition of ALK5 and p38alpha, and TGFβ reporter activity.

| | Kinase inhibition<br>IC<sub>50</sub> (nM) | | | Cellular inhibition TGFβ pathway<br>IC<sub>50</sub> (nM) | | |
| --- | --- | --- | --- | --- | --- | --- |
|  | HYL001 | Galunisertib | Vactosertib | HYL001 | Galunisertib | Vactosertib |
| ALK5<br>(TGFB1) | 177 | 448 | 51 | 72 | 157 | 10 |
| p38α | 548 | 920 | 6893 | - | - | - |

We next tested the three inhibitors in a cellular assay, using a luciferase-based reporter assay^20^ in HEK293T cells in the presence or absence of inhibitors (Table 2). Upon activation of the endogenous receptors by 5 ng/ml recombinant TGFβ, phosphorylation of the TGFBR1 triggers phosphorylation of the intracellular mediators SMAD2/3, and subsequent nuclear translocation with SMAD4. The SMAD2/3-SMAD4 complex binds to the TGFβ response element (12xCAGA) allowing the transcription of the Firefly luciferase reporter. In this assay, inhibition of TGFβ activity by HYL001 showed again an approximate 2-fold higher potency compared to galunisertib (Figure 5, Table 2). Vactosertib was more potent *in vitro* than HYL001 and galunisertib.

**Figure 5.**
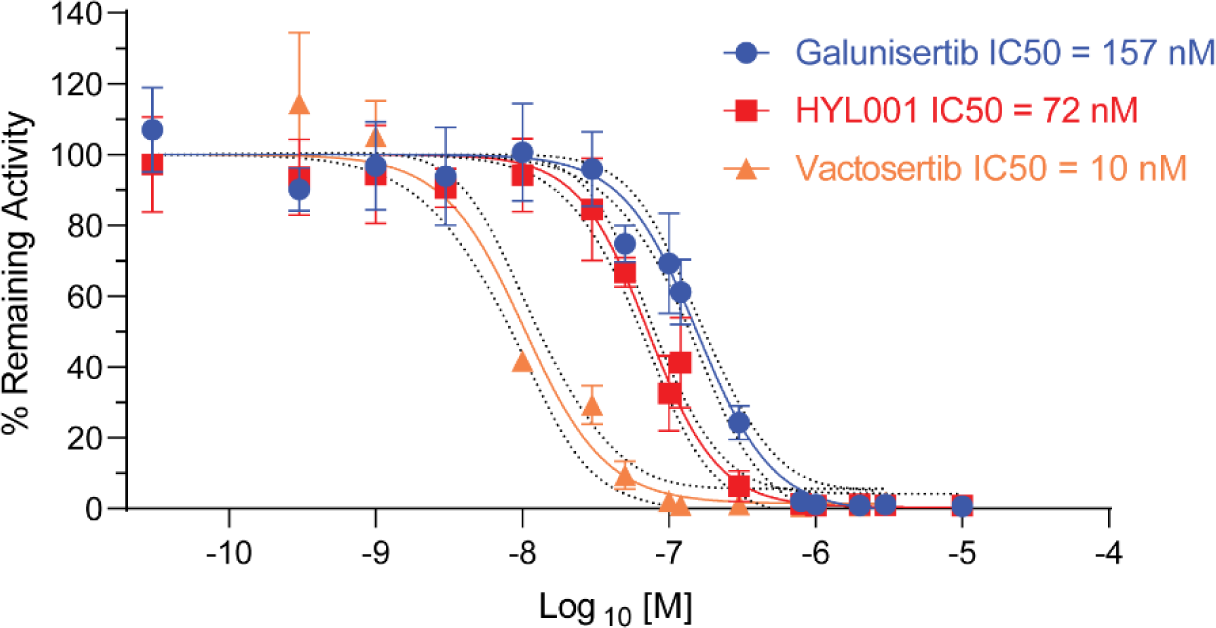
Cellular TGFβ reporter assay. Inhibition of TGFβ signaling in a cellular transcriptional reporter assay (0.03–10,000 nM compound). Data are means ± SD from 2–4 independent experiments. 95% CIs are indicated.

### Pharmacological characterization

To advance in the pre-clinical development of HYL001, we engaged in studies of its *in vitro* distribution and metabolism (DM) and *in vivo* pharmacokinetics (PK). The determination of the free fraction (unbound drug) of the drug is important for potential drug candidates. We evaluated the protein binding of HYL001 in human, rat and mouse plasma by equilibrium dialysis, a widely used method for protein binding measurements (Table 3). This was accomplished by spiking HYL001 at 5 µM into plasma and dialyzing against buffer until equilibrium was achieved (4 hours). Analyte area ratios of HYL001 in plasma and buffer were determined to calculate unbound and bound percentages of compounds to the plasma proteins. The plasma protein binding (PPB) of warfarin was analyzed in parallel as a standard reference. HYL001 showed high binding to human, rat, and mouse plasma proteins (Table 3).

**Table 3.** Plasma protein binding of HYL001 in 3 species (n= 3)

| Compounds | Species | % Bound<br>Mean ± SD | % Free<br>fraction | %<br>Recovery | % Compound<br>remaining at 4 hrs |
| --- | --- | --- | --- | --- | --- |
| **Warfarin** | Human | 99.1 ± 0.1 | 0.9 | 100.2 | 84.2 |
|  | Rat | 99.1 ± 0.1 | 0.9 | 98.6 | 104.8 |
|  | Mice | 97.2 ± 0.5 | 2.8 | 99.9 | 106.2 |
| **HYL001** | Human | 91.6 ± 1.1 | 8.4 | 100.6 | 101.1 |
|  | Rat | 95.1 ± 0.8 | 4.9 | 100.3 | 106.1 |
|  | Mice | 95.2 ± 0.3 | 4.8 | 112.4 | 113.1 |

Subsequently, we evaluated the metabolic stability of HYL001, galunisertib and vactosertib in human cryopreserved hepatocytes (Table 4). Primary hepatocytes contain the enzymes and co-factors needed for phase I and phase II drug metabolism, making them a well-established *in vitro* model for assessment of drug metabolic stability. In these assays, both HYL001 and vactosertib showed a similar pattern with <50% parent compound remaining in human hepatocytes within 1 hour of incubation (Table 4). Comparatively, galunisertib was more stable with 76% remaining after 1 h incubation. All three compounds were stable in buffer during the experiment. The results of the control compounds (testosterone and OH-coumarin) were consistent with historical data (not shown).

**Table 4.**
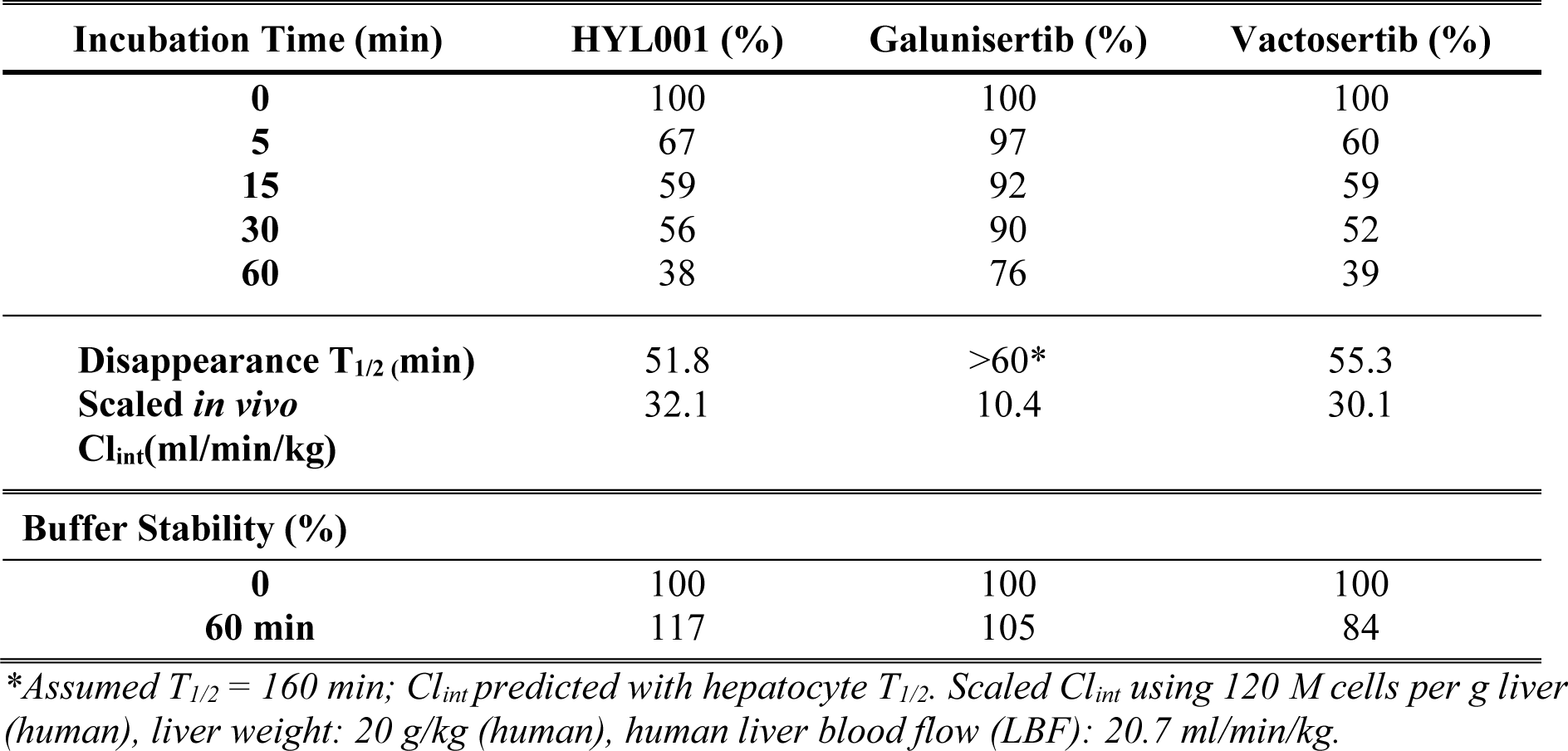
Metabolic stability assessed with human hepatocytes.

| Incubation Time (min) | HYL001 (%) | Galunisertib (%) | Vactosertib (%) |
| --- | --- | --- | --- |
| 0 | 100 | 100 | 100 |
| 5 | 67 | 97 | 60 |
| 15 | 59 | 92 | 59 |
| 30 | 56 | 90 | 52 |
| 60 | 38 | 76 | 39 |
| Disappearance $T_{1/2}$ (min) | 51.8 | >60* | 55.3 |
| Scaled *in vivo*<br>$Cl_{int}$(ml/min/kg) | 32.1 | 10.4 | 30.1 |
| **Buffer Stability (%)** |  |  |  |
| 0 | 100 | 100 | 100 |
| 60 min | 117 | 105 | 84 |

\*Assumed $T_{1/2} = 160$ min; $Cl_{int}$ predicted with hepatocyte $T_{1/2}$ . Scaled $Cl_{int}$ using 120 M cells per g liver (human), liver weight: 20 g/kg (human), human liver blood flow (LBF): 20.7 ml/min/kg.

We assessed the *in vivo* PK properties of HYL001 after a single oral administration (dose 71 mg/kg) in BALB/c mice as well as in Sprague Dawley Rats. Liquid chromatography–tandem mass spectrometry (LC/MS/MS) was used for the determination of HYL001 in plasma (Figure 6 and Table 5). Inspection of the plasma concentration−time profile for HYL001 revealed that the mean value of the maximum plasma concentration (C_max_) after oral dosing of 71 mg/kg was 2050 ±1459 ng/mL in mice, and 8542 ±1220 ng/mL in rat. Peak plasma concentrations were observed at 0.25 h (mice, first time point tested) and 0.67 ±0.29 h (rats), suggesting rapid absorption after oral dosing.

**Figure 6.**
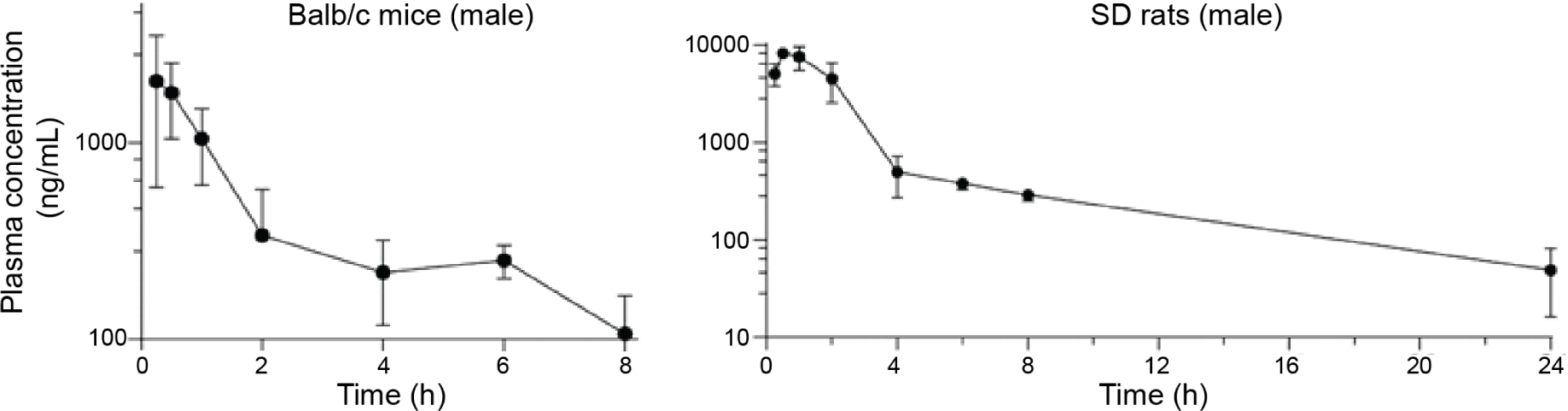
Plasma concentrations of HYL001. Longitudinal measurements after oral administration of a single dose (71 mg/kg) in Balb/c mice and Sprague Dawley (SD) rats.

**Table 5.** Pharmacokinetics of HYL001 in preclinical species after oral administration. Data shown for galunisertib^20^ and vactosertib^18,21^ are from indicated publications.

| | Species | Dose<br>(mg/Kg) | $T_{max}$<br>(h) | $C_{max}$<br>(ng/mL) | $AUC_{last}$<br>(ng.h/mL) | $AUC_{inf}$<br>(ng.h/mL) | $T_{1/2}$<br>(h) | $V_{ss}/F$<br>(L/kg) | $Cl/F$<br>(mL/min/kg) |
| --- | --- | --- | --- | --- | --- | --- | --- | --- | --- |
| **HYL001** | Mice<br>(♂ BALB/c) | 71 | 0.25 | 2050 | 3520 | 4110 | 2.45 | 66.7 | 288 |
| **Gal**<sup>20</sup> | (♀ CD-1) | 75 | 0.5 | 3330 | 3110 | NA | 1.4 | NA | 401.9 |
| **Vacto**<sup>21</sup> | (FVB/N) | 10 | 0.5 | 625 | 724 | 752 | 3.26 | 18 | 222 |
| **HYL001** | Rats<br>(♂ SD) | 71 | 0.67 | 8542 | 21575 | 22007 | 4.8 | 10.8 | 53.8 |
| **Vacto**<sup>18</sup> | (♀ SD) | 10 | NA | 1620 | NA | 1426 | 2.5 | NA | NA |

Coefficient of variation in mice ranged between 19–71%. $C_{max}$ : maximum plasma concentration; $T_{max}$ , time to $C_{max}$ ; $V_{ss}/F$ apparent volume of distribution at steady state; $F$ : bioavailability; $Cl/F$ : apparent total body clearance; Values in italics were estimated using mean plasma levels and NCA with Kinetica software v6.0. For Gal: $Cl/F$ estimated using $AUC_{last}$ <sup>20</sup>; NA: not available.

The area under the curve for plasma concentration−time, extrapolated to the last time point after oral dosing (AUC_last_) was 3520 ng·h/mL for mice and 21575 ng·h/mL for rats (Table 5). The elimination kinetics of HYL001 demonstrated a moderate terminal half-life (T_1/2_): approximately 2.5 and 5 h in mice and rats, respectively. These values are higher than those observed for galunisertib and vactosertib^18,20^.

As a caveat, small molecule PK may have sex differences in rodents because of differential metabolic enzymes (e.g.: CYP450). Having this in mind, HYL001 seems to show a lower clearance—and longer half-life—than vactosertib.

### Mutagenic potential

The mutagenic potential of HYL001 was evaluated by Ames test, assessing its ability to induce reverse mutations in the histidine operon of *Salmonella typhimurium* strains that allow for detection of both substitution mutations (TA100 and TA1535), and frameshift mutations (TA98 and TA1537). The mutagenic potential of HYL001 was tested at 6 concentrations (semi-log 1–320 μg/ml) alone or in the presence of liver S9 fractions. No positive mutagenic responses were observed with any of the strains (Table 6).

**Table 6.** Mutagenic potential assessed in *S. typhimurium* strains.

| *S. typhimurium* strain | Mutagenic data points |  | Dose dependency |  |
| --- | --- | --- | --- | --- |
|  | w/o S9 | with S9 | w/o S9 | with S9 |
| TA98 | No | No | No | No |
| TA100 | No | No | No | No |
| TA1535 | No | No | No | No |
| TA1537 | No | No | No | No |

### *In vitro* safety pharmacology

To identify potential adverse off-target drug interactions and de-risk the development of HYL001 as a drug^22^, we performed binding and enzymatic inhibition assays with a panel of key proteins (mainly cell receptors, neurotransmitter transporters and ion channels). Binding causing >50% inhibition—considered to represent a potential safety concern—was not observed to any of the targets for HYL001 (Figure 7A), nor did HYL001 cause >50% inhibition of any of the tested enzymatic activities (Figure 7B).

**Figure 7.**
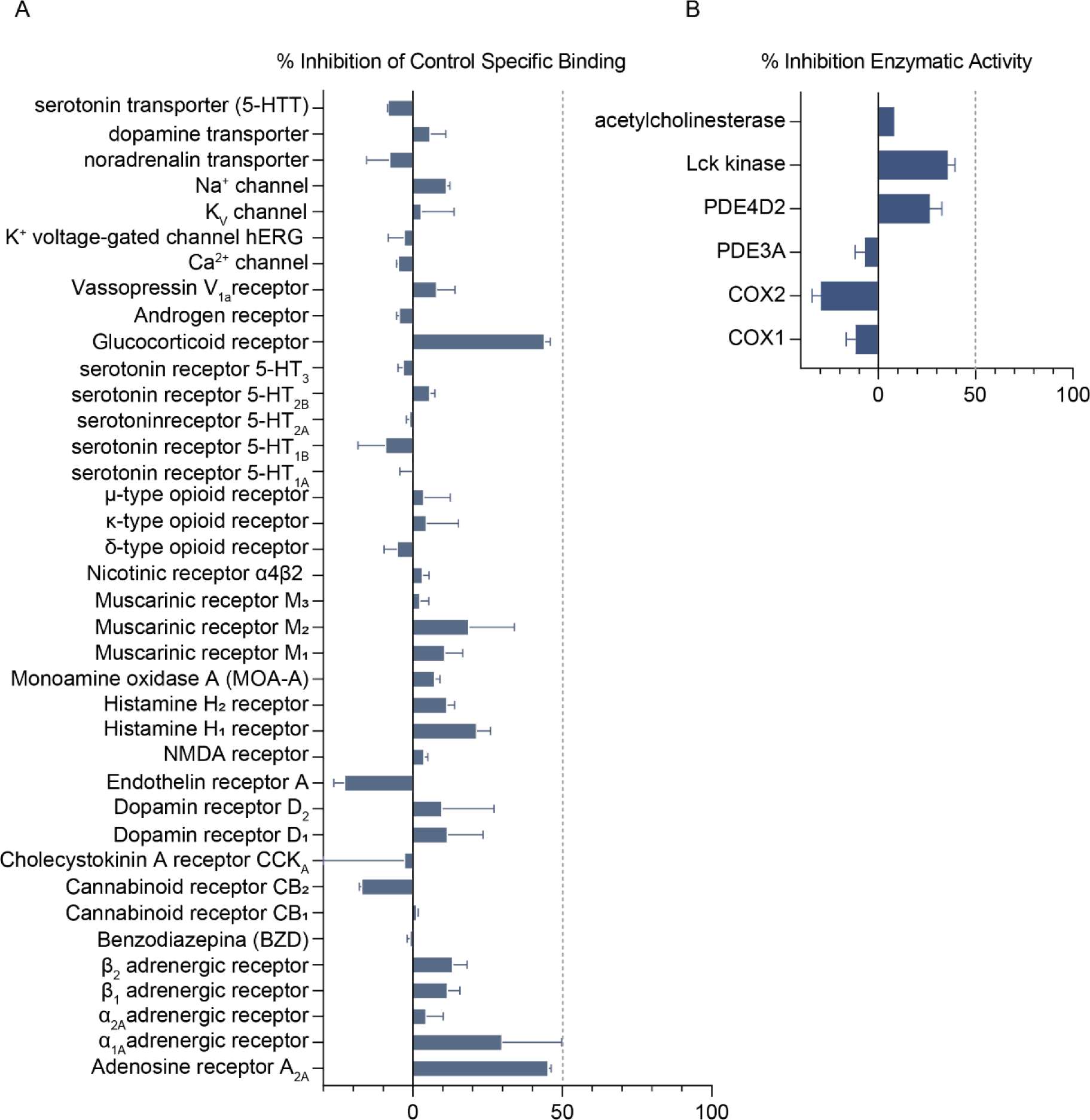
*In vitro* safety pharmacology. Binding (**A**) and enzymatic inhibition (**B)** of indicated receptors, protein channels and enzymes by 10 μM HYL001. HYL001 binding was calculated as the % inhibition of the binding of a radioactively labeled ligand specific for each target. HYL001 enzyme inhibition effect was calculated as % inhibition of control enzyme activity. Values are averages of two measurements ±SD.

Of note, inhibitory binding of the human Ether-à-go-go-Related Gene (hERG) potassium channel—which is not the case for HYL001 (Figure 7A)—is indicative of cardiotoxicity. Given the history of TGFβ inhibitors with on-target toxicity^14^, we further determined the activity of HYL001 towards hERG more specifically using a patch-clamp assay^23^. Unlike reference compound, E-4031 that blocks potassium channels of the hERG-type (exhibiting an IC_50_ of 37 nM, data not shown), HYL001 showed low inhibition at micromolar concentrations (Table 7). For comparison, the IC_50_ of vactosertib towards the hERG channel is 31.04 μM^18^, indicating a lower cardiac toxicity risk for HYL001.

**Table 7.** hERG inhibition cardiac toxicity assessment for HYL001.

| Concentration (μM) | Inhibition (%) |
| --- | --- |
| 3.7 | 1.8 |
| 11.1 | 8.4 |
| 33.3 | 22.8 |

### Pre-clinical toxicity in rodents

We performed exploratory pre-clinical toxicology studies with HYL001 and vactosertib in Sprague Dawley (SD) rats and C57BL/6J mice. Vactosertib, an ALK5 inhibitor of similar potency that is being tested in patients, was included as comparator drug.

#### Toxicology in SD rats

Four pre-clinical exploratory toxicology studies were conducted in rats (n = 6; 3 females + 3 males in all studies; the same numbers of vehicle control-treated rats were included, in which no abnormalities were detected). These were (group 1) a 4-week toxicity study with HYL001 at continuous dosing with 25 mg/kg twice daily (*bis in die*, BID), and (group 2) the same with a dose of 71 mg/kg BID, (group 3) the latter dose for 4 weeks, followed by a 28-day recovery period. The 4^th^ group received vactosertib for 4 weeks at the continuous dose of 82 mg/kg BID without recovery (equimolar with 71 mg/kg BID HYL001, similar to group 2; Figure 8).

**Figure 8.**
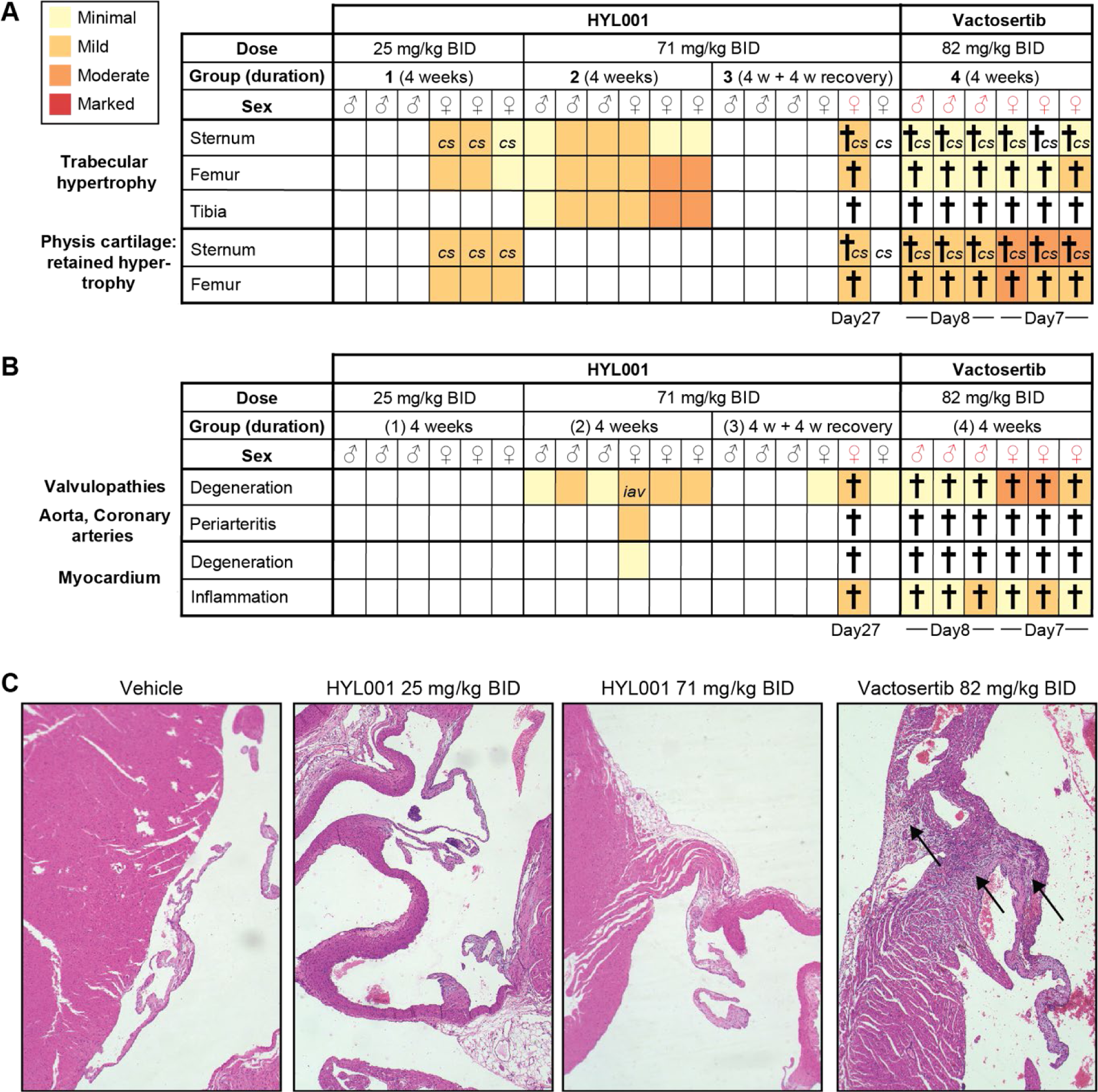
Rat toxicology study. (**A–B**) Overview of histopathological findings in bones (A) and the heart (B) for four groups of rats with sex and (planned) duration indicated, as well as occurrence and time of animal death/clinical endpoint (†). The colour scheme represents a 4-tier scale. Observations of curved sternum (*cs*) and chronic inflammation of the aortic valve (*iav*) are marked. (**C**) Examples of rat heart valves and adjacent myocardium in a healthy control, or after indicated treatment. Arrows indicate degeneration of the valve and chronic inflammation of adjacent myocardium. All images at 10x magnification.

There were no HYL001-related effects on body weight (gain), feed consumption, hematology, clinical chemistry analytes in rats of both sexes treated orally with HYL001 at either dose level. Also, no clinical findings were noted during treatment at 25 mg/kg/ BID. However, 3/6 female animals treated with 71 mg/kg BID (all from group 3) developed signs of lethargy, tachycardia and protrusion of sternum from day 8 of treatment. One of these females died on day 27 (just before recovery). The 2 remaining females from this group recovered from lethargy during the off-treatment period. Males in group 3 (71 mg/kg BID HYL001) only showed rough hair coats during the last two weeks of treatment, from which they soon recovered. 17/18 (94%) rats treated with HYL001 survived until end of study.

In contrast, in vactosertib-treated rats (group 4), animals gained signs of lethargy, protruded sternums and mild reduced locomotor activity (in males); and of lethargy, tachycardia, mild reduced locomotor activity, rough coat, piloerection and protruded sternum (in females), all within the first week of treatment. Strikingly, all female rats were found dead on day 7 and all male rats had to be sacrificed on day 8 (Figure 8A).

Given the previously reported on-target side-effects of TGFβ inhibitors^14,18,24,25^, detailed histopathological analysis focused on bones, heart, and gastrointestinal system was carried out in all groups.

##### Bones

Some female rats (HYL001) and all vactosertib-treated animals showed a curved sternum at death or end point. Increased bone formation in trabeculae (trabecular hypertrophy) was observed in femur, tibia and sternum, in both sexes and with both compounds (Figure 8A). This effect appeared to be reversible, as it was not found after the 4-week recovery period in group 3. Retained hypertrophied cartilage physis was identified in sternum and femur of all vactosertib-treated animals (mild–moderate) and in a few female HYL001-treated animals (mild). The latter abnormalities have been described before for ALK5 inhibitors^13,25^ yet are predicted to have low impact in adult humans, since their growth plates are expected to be closed. Also, these effects did not prevent galunisertib or vactosertib entering clinical trials.

##### Heart

The major safety concern regarding ALK5 inhibitors is the on-target cardiac toxicity observed in prior pre-clinical studies, which included valvulop athies (degenerative and inflammatory valvular lesions), myocardial degeneration/necrosis, hemorrhage, or mixed cell infiltrates in the myocardium, necrosis with inflammation of coronary arteries, and mixed cell infiltrates in the atrium^14,18,26^. We observed minimal to mild degeneration of heart valves in both male and female HYL001-treated rats at terminal examination after 4 weeks of repeated dosing in group 2 as well as the female from group 3 that died on-treatment (Figure 8B, C). This consisted of myxomatous expansion of the stroma or stromal proliferation. However, the heart valves of HYL001-treated rats were almost normal after a 4-week recovery. Relatedly, an increase in absolute and relative weight of the heart was found in female HYL001-treated rats after 4 weeks treatment (group 2), yet this was much reduced after 4 weeks of recovery (group 3). With vactosertib, moderate degeneration of valves was already observed after 7–8 days of treatment, when the rats died or had to be sacrificed. Both the curved sternum and increased weight of heart have been reported previously for galunisertib^24^ that was later safely used in patients. Furthermore, vactosertib is also being used in patients.

Inflammation of the aortic valve and periarteritis in aorta and coronary arteries were found in one female rat under HYL001 treatment (group 2), yet in none of the rats of group 3 (recovery group). These abnormalities were not observed in vactosertib treated animals at their (unintendedly) early endpoints. However, at that time, chronic inflammation of the myocardium was observed in all 6 vactosertib treated animals, as well as in the female rat treated with HYL001 that was found dead (Figure 8B, C). Only one female rat in group 2 showed minimal degeneration of myocardium, characterized by infiltration of mononuclear cells (predominantly lymphocytes and macrophages) and necrosis of cardiomyocytes. None of these were observed in the third group (recovery group) (Figure 8B). In fact, cardiac toxicity observations were absent or minimal in the 5 remaining rats of this recovery group, suggesting reversibility. In addition, no abnormalities in the heart were observed in rats from either sex in low-dose group 1 (Figure 8B). In this group, mild histopathological findings were observed in the sternum and femur of female rats only. As mentioned above, these are predicted to have a low impact in adult patient populations. For male rates, the no-observed-adverse-effect-level of HYL001 was established at the dose of 25 mg/kg BID, similar to the reported NOAEL of vactosertib^18^.

##### Gastrointestinal tract

No microscopic findings were observed along the gastrointestinal tract (stomach, duodenum, jejunum, ileum, caecum, colon and rectum), except minimal hyperplasia of the non-glandular stomach and chronic inflammation observed in one female rat under vactosertib treatment.

#### Toxicology in Balb/C mice

In comparison to rats, toxicity of TGFβ receptor-1 inhibitors in Balb/c mice was less severe (Figure 9). No mortality or signs of morbidity were observed for mice treated with HYL001 or vactosertib at equimolar dose (71 mg/kg BID and 82 mg/kg BID, respectively). Specifically, there were no compound-related effects on body weight, feed consumption, hematology, and clinical chemistry analytes in mice of both sexes, except for a slight increase in neutrophil count for vactosertib-treated males. Histopathology examination in bones found relatively mild phenotypes (Figure 9A).

**Figure 9.**
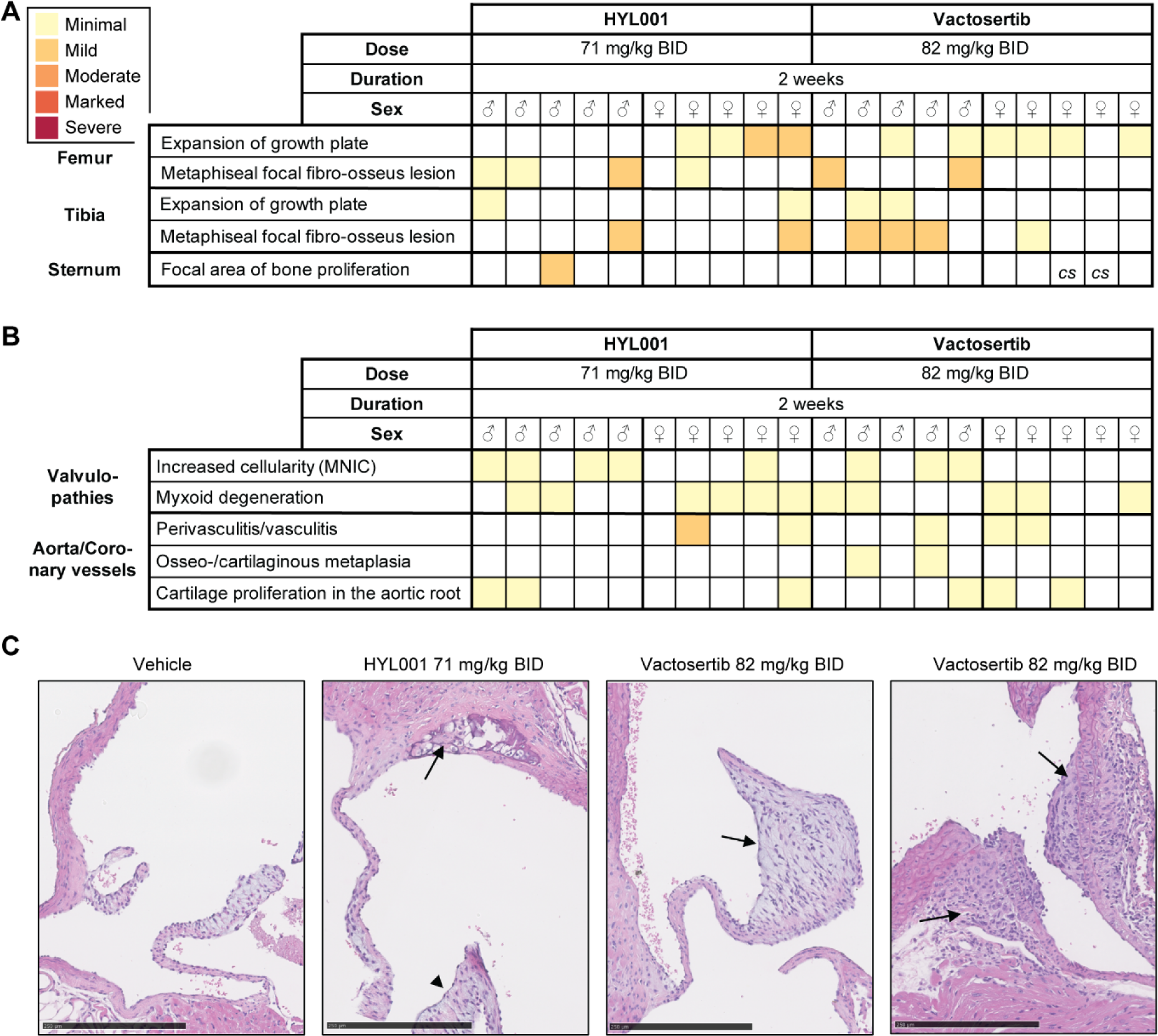
Mouse toxicology study. (**A–B**) Overview of histopathological findings in bones (A) and the heart (B) for indicated sex and treatment. The colour scheme represents a 5-tier scale. Observations of curved sternum (*cs*) are indicated. *MNIC:* mononuclear immune cell. (**C**) Representative examples of a semilunar valve from vehicle control-treated mouse (left); a semilunar valve with myxoid degeneration (arrowhead) and cartilaginous metaplasia in the root of aorta in a HYL001-treated animal (arrow; middle left); a semilunar valve with myxoid degeneration and increase in cellularity from vactosertib treated mouse (arrow; middle right); and focal mononuclear perivasculitis/vasculitis in coronary vessels in a vactosertib treated mouse (arrow, right). All images at 20X magnification.

##### Bones

Minimal–mild signs of proliferation of fibro-osseous tissue in the metaphysis (bone hypertrophy) were observed in mice with either compound, especially in femur and tibia. Despite two cases of macroscopically curved sternum, histopathological observations were almost lacking there (Figure 9A).

##### Heart

There were no alterations in any of the myocardia. However, similar to the rats, treated mice exhibited some minimal to mild myxoid degeneration of heart valves—which were also observed in 2/10 vehicle control-treated mice (not shown)—or an increase in cellularity mainly from mononuclear immune cells (Figure 9B, C). Mice under both treatments presented focal mononuclear perivasculitis/vasculitis in coronary vessels. Some mice under both treatments also showed associated incipient osseous/cartilaginous metaplasia at the root of the aorta or coronary vessels. This phenotype was more pronounced in vactosertib treated mice.

##### Gastrointestinal tract

No observations were made, except for slight inflammatory infiltrates, mainly with mononuclear cells, in some female livers (2/5 mice treated with HYL001 and 4/5 for vactosertib).

Taken together, female animals exhibited more treatment-related toxicities than males, and rats were more sensitive than mice. Importantly, HYL001 toxicity in SD rats appears to be milder than that of an equimolarly dosed compound currently in clinical trials.

### *In vivo* HYL001 efficacy as immunotherapy against CRC metastases

We next assessed the *in vivo* efficacy of HYL001 in our model of pMMR/MSS CRC^5^. As reported, we generated a genetic mouse model bearing mutations in key CRC pathways (WNT, P53, MAPK and TGFβ) and derived mouse tumour organoids (MTOs) from invasive adenocarcinomas as well as liver metastases generated in quadruple compound mice. Both the genetic model and mice implanted with MTOs developed intestinal tumors and metastatic liver lesions that reproduced several key features of human poor prognosis CMS4-like CRC, including a stroma-rich, TGFβ-activated TME and low levels of T cell infiltrations^5^. Subsequently, this MTO-based system has proven a powerful pre-clinical model, both to dissect epithelial tumour cell heterogeneity that is representative for human cancer cell states^27,28^ and to further dissect the TME for additional mechanisms of (dys)regulation of cancer immunity^29–32^.

We first assayed the capacity of HYL001 to inhibit liver metastasis formation as monotherapy. In this model, we inoculated MTOs derived from liver metastases into the portal vein (by intrasplenic or direct intravenous injection) to model their becoming trapped in hepatic sinusoids and, depending on their metastatic capacity, initiation of liver nodules. We had previously demonstrated that galunisertib can prevent the formation of liver metastases using this model, albeit at a very high dose (720–800 mg/kg BID)^5^, which is likely far above clinically relevant levels. Normalizing for the 800 mg/kg BID galunisertib dose, we tested a dose range of 0.03×–0.9× molar equivalents (21–637 mg/kg BID) of HYL001. Remarkably, a 0.1× molar equivalent (71 mg/kg BID) of HYL001 was sufficient to prevent metastasis formation in 10 out of 11 animals. All treated animals developed metastases with galunisertib at this 0.1× dose level (80 mg/kg BID) (Figure 10).

**Figure 10.**
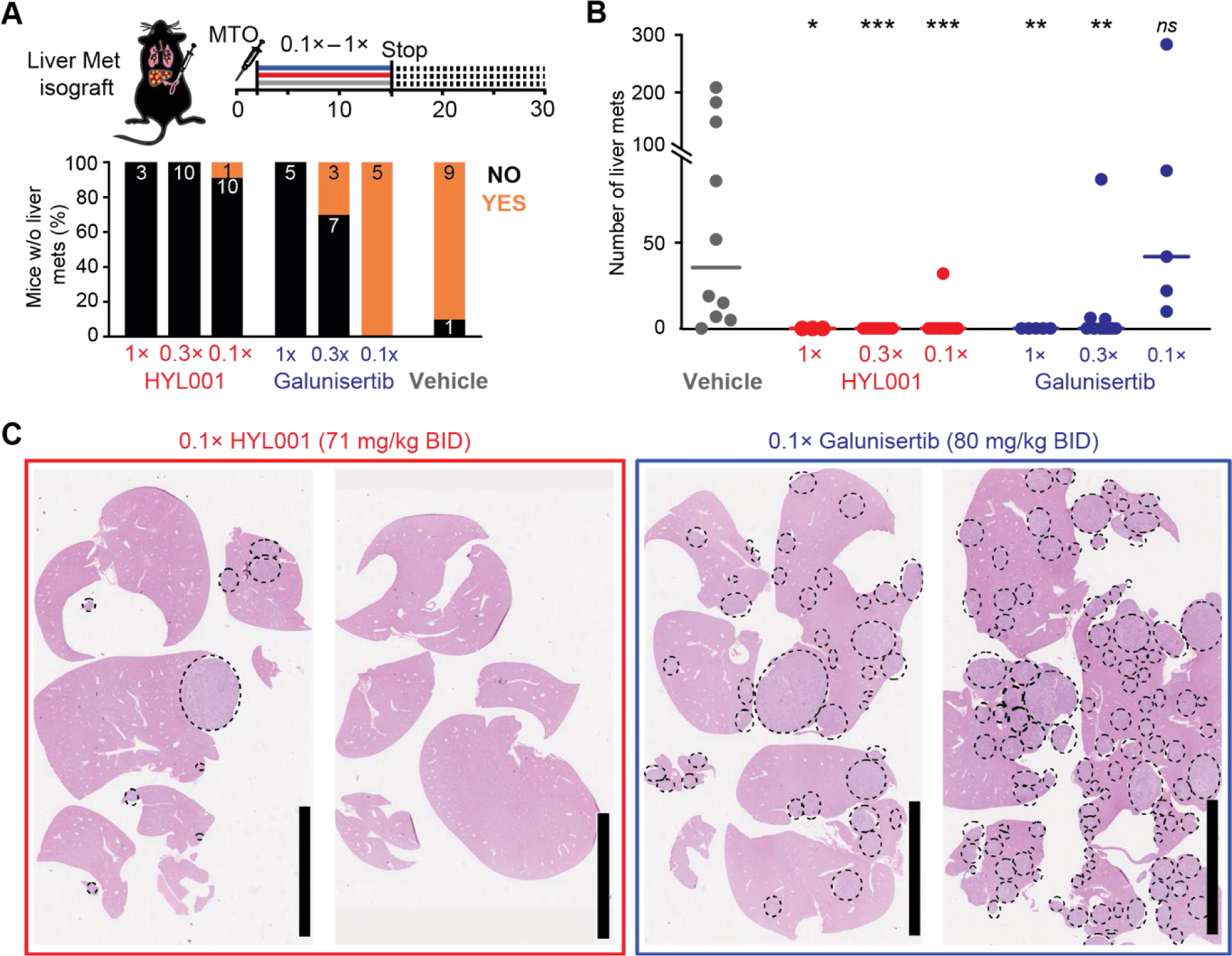
HYL001 prevents metastasis formation *in vivo*. (**A**) Experimental scheme, including injection of MTOs. Experimental endpoint was at day 28, two weeks after treatment stop. Below: the number of mice treated that had (in orange) or did not have (in black) detectable liver metastases.^5^ (**B**) Liver metastases (Mets) were counted at endpoints; each point represents one mouse. Non-parametric Mann-Whitney U test, two-tailed. ns: P > 0.05; * P < 0.05; ** P < 0.01; *** P < 0.001; (**C**) Representative images of H&E whole slides containing liver sections from mice treated with TGFβ inhibitors; dashed line circles delimit metastases. 1x, efficacious dose for galunisertib established in ref. 5: 800 mg/kg BID, and molar equivalent for HYL001 (710 mg/kg BID). 0.3x and 0.1x: dilutions thereof. Bars, 10 mm

Moreover, treatment of mice that had already established liver metastases with HYL001 resulted in a marked decrease in protein expression of relevant TGFβ targets, including phospho-SMAD2, CTHRC1, and Caldesmon^9^ (Figure 11).

**Figure 11.**
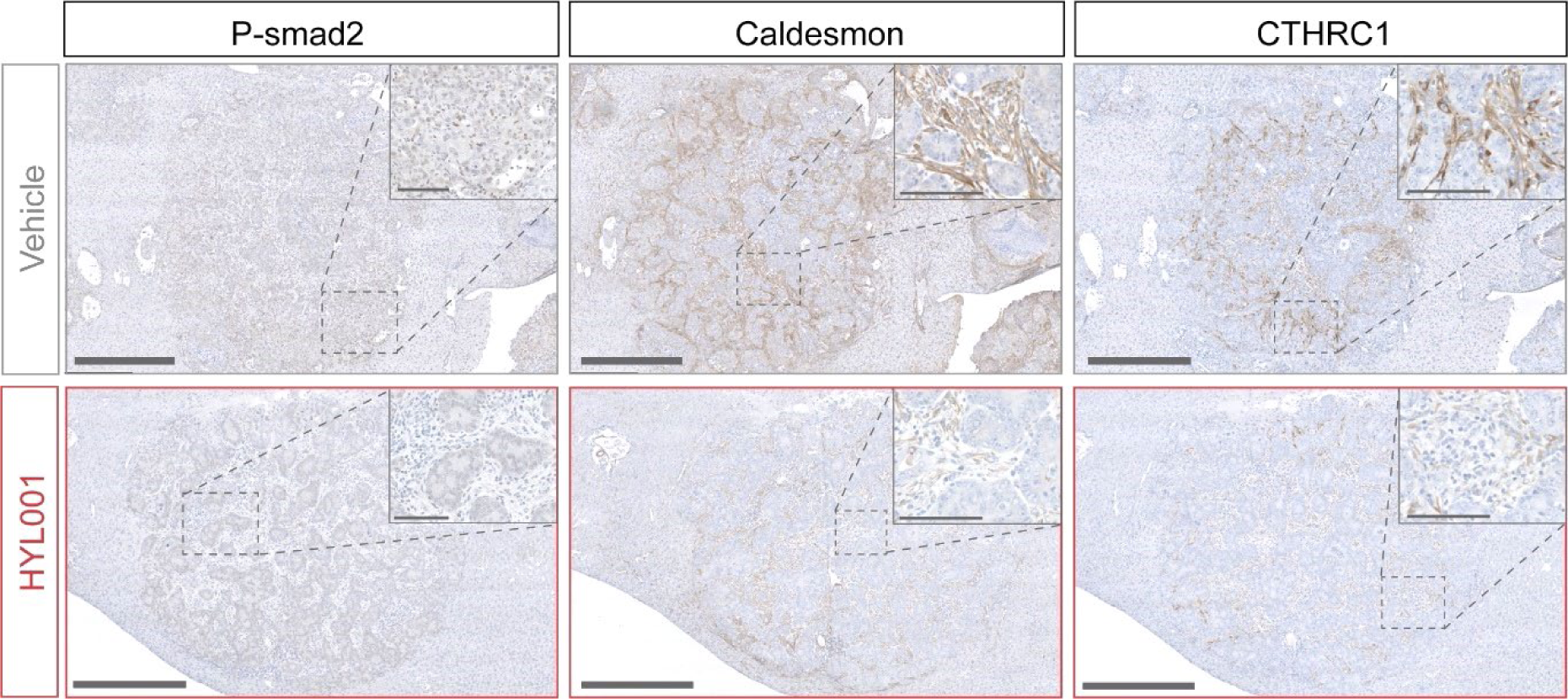
HYL001 treatment reduces protein levels of TGFβ signalling targets *in vivo*. Representative images of the indicated protein expression levels assessed by immunohistochemistry in liver metastases of mice with established liver metastases treated for 3 days with HYL001 at 0.3x dose. Bars 500 μm in 10x magnification images; 100 μm in insets.

Despite the role of TGFβ blockade in preventing liver metastatic initiation in our mouse model, which we showed to involve T cell-mediated anti-cancer immunity^5^, established metastatic CRC was not cured by this treatment in our model. Moreover, these mice were resistant to immune checkpoint therapy like human patients with pMMR/MSS CRC are in the clinic^5,33^. However, we showed that TGFβ inhibition mediated by galunisertib (at high concentrations) combined with immune checkpoint therapy could overcome such resistance and synergize with immunotherapies to cure established metastases^5^. For these experiments, we used the 1x galunisertib dosage (800 mg/kg BID)^5^. Encouraged by the increased potency observed *in* vitro and in the inhibition of liver metastatic initiation *in vivo*, we evaluated the ability of a relatively low HYL001 dose to help overcome established liver metastases, when combined with immune checkpoint inhibition. In this experimental set up, a low number of MTO cells (25K) was injected in the portal vein (leaving the spleen unaltered) and liver metastases were allowed to develop and grow for 15 days, before treatment with TGFβ inhibitors was initiated. This was done in combination with 3–4 intraperitoneal (IP) injections of immune-checkpoint inhibition therapy (anti-PD1 or isotype control antibody), spaced 2–3 days (Figure 12A, B). In accordance with our previous results^5^, immune checkpoint therapy alone was not efficacious to cure liver established metastases, and neither were the three TGFβ inhibitors as monotherapy (80, 82 and 71 mg/kg BID equimolar 0.1x doses for galunisertib, vactosertib and HYL001, respectively; Figure 12A). In dual therapy, HYL001 exhibited highest efficacy curing up to 80% mice (8/10) at endpoint (surviving for 2.5 months after injection, the study endpoint), and a marked overall reduction of tumour burden in numbers of liver nodules (Figure 12A, B).

**Figure 12.**
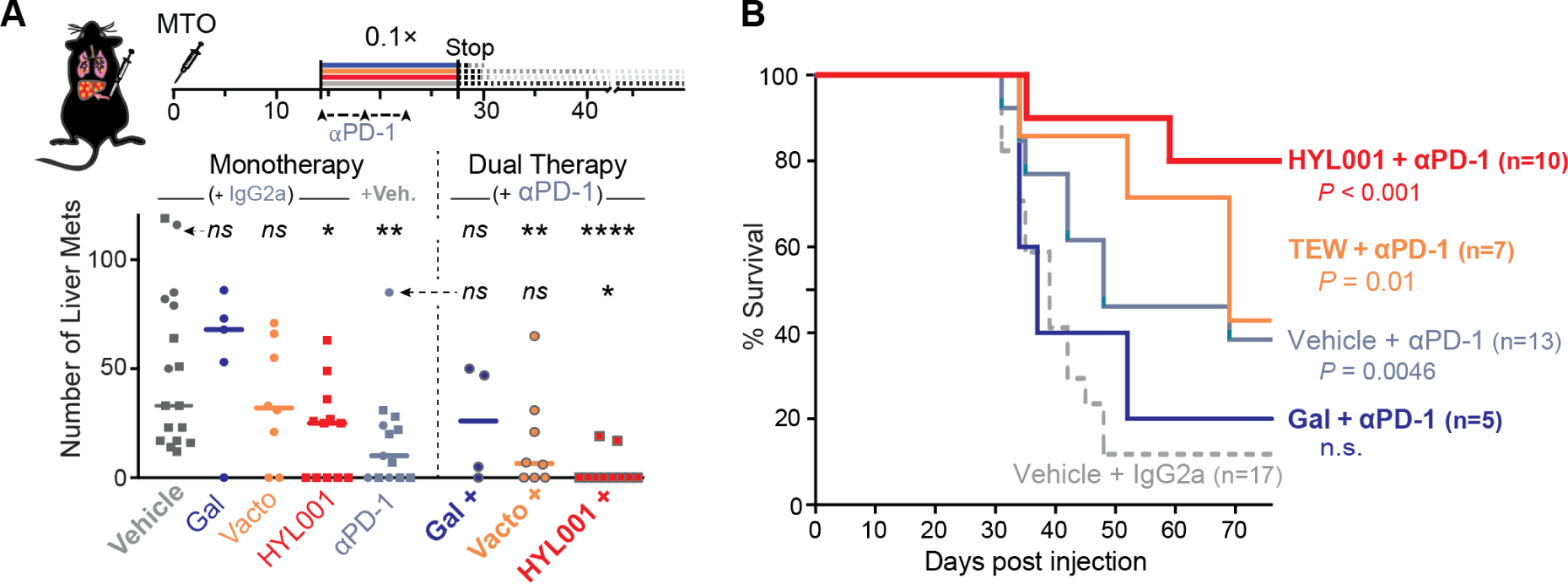
HYL001 efficacy in dual immunotherapy against established CRC liver metastases. **A**) Overview of experimental timing: dual treatment between day 14–28 after MTO injection (TGFβ inhibitor doses were all 0.1×); follow-up until humane endpoint. Below: numbers of countable liver lesions after sacrifice. Statistical comparisons to the vehicle+IgG2a control (above) or the vehicle+αPD1 condition (below) by two-tailed Mann-Whitney U test; *ns*: P ≥ 0.05; * P < 0.05; ** P < 0.01; *** P < 0.001; **** P < 0.0001. **B**) Kaplan-Meier curves indicate the survival (until humane endpoint) of mice under each treatment; *P* values were computed by Mantel-Cox test (Log-rank).

## DISCUSSION AND CONCLUSIONS

Despite long-standing and active interest in therapeutically targeting the TGFβ pathway, drug development has not yet led to successful treatments due to low clinical efficacy and/or high toxicity in humans^6,13,34^. Here we present HYL001, a TGFBR1/ALK5 inhibitor that has increased *in vitro* and *in vivo* potency at relatively low doses and has favourable preclinical safety characteristics. We show that HYL001, while structurally related to galunisertib, is functionally superior in eliciting T cell-mediated eradication of CRC liver metastasis in mice. In direct comparison with vactosertib (TEW-7197), an unrelated second-generation inhibitor that is in clinical trials for solid cancers including CRC, HYL001 has similar or better efficacy at an equimolar dose range in our mouse experiments, albeit a safer toxicity profile in mice and rats. Therefore, HYL001 is a promising inhibitor of TGFβ signalling with clear preclinical benefit—offering an economical advantage for *in vivo* research—that also warrants clinical testing.

Whereas the increased efficacy of HYL001 may arise solely from a distinction in on-target affinity due to the structural difference with galunisertib, the higher degree of improvement *in vivo* over *in vitro* (∼9-fold *vs* ∼2.5-fold) suggests an additional effect. We investigated plasma protein binding, hepatic stability and PK in this study. The reduced metabolic stability in hepatocyte culture, compared to galunisertib, may be related to the relatively low (∼91%) plasma protein binding value for HYL001 in humans—suggesting a potentially higher unbound fraction. On the other hand, this may also be associated to elevated drug–target interaction—although *in vivo* drug kinetics are rather more complex than can be captured by a PPB value^35^. Towards a more comprehensive understanding thereof, our PK studies show good gastrointestinal absorption and indicate clearance values for HYL001 that are slower than both galunisertib and vactosertib. Therefore, the increased efficacy in vivo might be attributed to the sum of favorable traits in a number of variables.

Measuring *in vitro* affinity for ALK5, vactosertib showed the lowest IC_50_, galunisertib the highest, and HYL001 has an intermediate value. We subsequently performed a selectivity screen using an *in vitro* competitive binding assay, and observed binding preference, besides for ALK5, towards ALK4/ACVR1B for all 4 compounds. We further found that galunisertib and HYL001 are moderately selective with only few differences—with HYL001 additionally showing >85% competitive binding to ABL-1, CSNK1G2, and p38α in our assay. Vactosertib is clearly less selective, showing >85% competitive binding to an additional set of targets that includes ERBB2, FGFR2/3, KIT, MEK1/2, PDGFRA/B, SRC, and VEGFR2. Whereas these distinct on- and off-target affinity profiles might translate into varying *in vivo* efficacy, we did not observe strong differences there. As poor selectivity for multitarget kinases has been hypothesized to lead to a concentration-dependent broadening of the inhibitory spectrum^36^, the lack of evident differences together with the relatively low doses used in our current study indicate that the immuno-oncological effect of inhibitors is indeed on-target.

Nevertheless, it is possible that the lower selectivity for vactosertib may be related to its comparatively elevated toxicity. Vactosertib is being given to patients with cancer at a 200– 600 mg daily dose^37–39^. Assuming an average weight of 60 kg, this would represent a 3–10 mg/kg dose, which in turn can be converted (using a body weight–surface area scaling ratio of 12.3)^40^ to an equivalent of a 41–123 mg/kg daily dosage in mice. This is just below to what we tested here and our 0.1× dose (164 mg daily—or a human equivalent of 800 mg) would therefore be expected to risk cardiac or other toxicities. Clinical data however suggest that these levels are sufficiently safe^37–39^, suggesting differences between toxicology findings in rodents and in patients. Indeed, we have shown relevant differences in toxicity also exist between rodents. In comparison, galunisertib required a ∼9-fold higher dosing range (in human equivalents >7 g daily) to be efficacious in our CRC model—dose levels that are unfeasible for human translation. The fact that the maximum-tolerated dose used in clinical trials is lower than what appears to be required for galunisertib may explain the lack of therapeutic window for this first-generation compound^13^. HYL001 at the dose levels used here (2 x 71 mg daily in mice; human equivalent: 692 mg/day) performed very well, with an even more favourable toxicity profile than vactosertib in either rodent model.

In our preclinical efficacy study, we used mouse tumour organoids introduced into portal vein circulation of immunocompetent (syngeneic) mice to induce the formation of liver metastases^5^. We described the benefit of using organoids over available cell lines previously and linked the characterization of our model—as mismatch-repair proficient, microsatellite-stable invasive adenocarcinoma with human-like histopathology—to the majority of cases of mCRC^5^. These patients are not likely to respond to immune checkpoint inhibitor monotherapy^41^, but may benefit from combinatorial immunotherapies such as the added blockade of TGFβ signalling^3,5,6^. Indeed, mounting evidence connects this pathway to dominant changes in the TME and suppression of immunity in particular^4^, yet they simultaneously emphasize the contextual, pleiotropic functions of TGFβ—some of which are linked to severe toxicities in the context of systemic inhibition. Besides leading to several alternative therapeutic strategies—including blockade of upstream activation, targeted ligand traps, and singling out individual isoforms^6,7,13^—there remains potential for a small molecule TGFβ receptor kinase inhibitor with relatively low toxicity and retained potency. In comparison to e.g. fusion protein TGFB ligand traps, having a 1:1 stoichiometry and high production cost, a compound (with a chemical handle) can be fine-tuned to enhance efficacy and tolerability, and should be more cost-effective. Moreover, preclinical data for some upcoming inhibitors (increasingly biologicals, e.g. SRK-181, bintrafusp alfa, …) often exerted only partial therapeutic responses in experimental models^26,42^, while effectively depleting plasma TGFB levels^43^—potentially indicating trap quenching before reaching the TME.

In conclusion, we have synthesized a new small molecular inhibitor targeting TGFBR1 activity that has the required potency to inhibit a TGFβ-rich stroma and synergize with immunotherapies in advanced models of poor prognosis CRC. HYL001 provides superior potency compared to galunisertib at 9-fold lower dosing, without added toxicity, and compares favourably to vactosertib, another second-generation compound. The combined data we present here support a clinical study to confirm safety and feasibility for prolonged and or dosing in patients. Subsequently, HYL001 or functional derivatives—taking advantage it its ‘handle’, should be tested to treat metastatic cancers that thrive in a highly TGFβ-mediated immunosuppressive microenvironment.

## EXPERIMENTAL SECTION

### Synthesis of compounds

All new compounds were chemically synthesized, purified by chromatography and characterized by ^1^H NMR, ^13^C NMR, IR and HRMS. Compounds were of a purity ≥95% as determined by HRMS and ^13^C NMR spectroscopy, or by HPLC/Ms. NMR spectra were recorded at 23°C on a Varian Mercury 400 or Varian 500 apparatus. ^1^H NMR and ^13^C NMR spectra were referenced either to relative internal TMS or to residual solvent peaks. IR spectra were recorded in a Thermo Nicolet Nexus FT-IR apparatus. Melting points were determined using a Büchi M-540 apparatus. HRMS were recorded in a LTQ-FT Ultra (Thermo Scientific) using nanoelectrospray technique. HPLC chromatography was performed on Hewlett-Packard 1050 equipment with UV detection using a Kinetix EVO C18 50x 4.6 mm, 2.6 μm column (Standard gradient: 10 mM NH_4_CO_3_ / MeCN (95:5) – (0:100)). All reactions were carried out under an inert atmosphere (N_2_) unless otherwise stated.

The following compounds were prepared following standard or reported procedures: galunisertib^8,44^, vactosertib^18,45^, 6-methoxy-4-methylquinoline **(3a)**^46,47^, 6-bromo-4-methylquinoline **(3b)**^16^, and 1-aminopyrrolidin-2-one tosylate salt^44^.

### 2-(6-methoxyquinolin-4-yl)-1-(6-methylpyridin-2-yl)ethan-1-one (4a)

A solution of methyl 6-methylpicolinate (5.05 g, 33.4 mmol) in 15 mL of dry THF was stirred under N_2_ atm. for 1h in the presence of 0.5 g of 3Å molecular sieves.

6-Methoxy-4-methylquinoline (**3a,** previously dried under Dean-Stark conditions; 2.90 g, 16.7 mmol) were dissolved in 20 mL of anh THF under N_2_ atm and the resulting solution was cooled to –40 °C. A 2M solution of LDA in THF (19.2 mL, 38.4 mmol) was added dropwise over the quinoline solution and the resulting dark green suspension was stirred for 30 minutes at –40 °C. After this time, the anion and the previously prepared ester solution were cooled down to –78 °C and the anion was added over the ester. The resulting mixture was left to achieve room temperature and stir overnight. Then, 100 mL of a saturated solution of NH_4_Cl was added and the aqueous layer was extracted with EtOAc (3×100 mL). The combined organic extracts were dried over anh MgSO_4_, concentrated *in vacuo* and purified by column chromatography (eluted with a gradient of hexanes and EtOAc from 100:0 to 0:100). The desired product (**4a**) was isolated as an orange oil (3.37 g, 69%). The spectroscopic properties were identical to the ones described^48^; ^1^H NMR (400 MHz, CDCl_3_) δ 8.70 (m, 1H), 8.04 – 7.95 (m, 1H), 7.84 (d, *J* = 7.6 Hz, 1H), 7.71 (t, *J* = 7.7 Hz, 1H), 7.43 – 7.31 (m, 4H), 4.95 (s, 2H), 3.83 (s, 3H), 2.67 (s, 3H) ppm. ^13^C NMR (101 MHz, CDCl_3_) δ 198.2, 158.1, 157.8, 152.2, 147.5, 144.5, 140.4, 137.2, 131.5, 128.9, 127.2, 123.8, 121.5, 119.6, 102.5, 55.4, 40.8, 24.4 ppm. IR (film): 3377, 3065, 2969, 2932, 2827, 1693, 1624, 1577, 1520, 1242, 1230 cm^-1^. HRMS (ESI): m/z [M + H]^+^ calculated for C_18_H_16_N_2_O_2_: 292.1212; found: 292.1208.

### 2-(6-bromoquinolin-4-yl)-1-(6-methylpyridin-2-yl)ethan-1-one (4b)

6-Bromo-4-methylquinoline (**3b**, 18.1 g, 81.7 mmol) was charged in a flask and purged with N_2_. Anh THF (345 mL) was added and the resulting solution was cooled to –20 °C. To this solution, NaHMDS 2M (123 mL, 245 mmol) was slowly added. The resulting mixture was stirred for 3h at –20 °C to form the corresponding anion. Under anhydrous conditions, a solution of methyl-6-methylpicolinate (13.3 g, 98 mmol) in anh THF (40 mL) was added dropwise to the anion solution *via cannula*. The resulting mixture was stirred at –20 °C for 18h. After completion, THF was concentrated until 10% of the initial volume was left. EtOAc (300 mL) and a saturated solution of NH_4_Cl (600 mL) were added and the biphasic mixture was stirred until all the solid was dissolved. Phases were separated and the aqueous phase was extracted with more EtOAc (2×300 mL). The resulting organic extracts were dried over MgSO_4_ and filtered, and solvent was removed under reduced pressure.

MeOH (675 mL) was added and the mixture was stirred at rt 16h. The resulting suspension was filtered and the yellow residue was rinsed with more MeOH. Solvent was removed under reduced pressure to afford a red solid (24.2 g, 87%) that was directly used in the next reaction. The spectroscopic properties were identical to the ones described^16^; ^1^H NMR (400 MHz, CDCl_3_) δ 8.86 (d, *J* = 4.6 Hz, 1H), 8.41 (d, *J* = 2.2 Hz, 1H), 8.15 – 8.08 (m, 1H), 7.90 – 7.85 (m, 1H), 7.81 (dd, *J* = 9.0, 2.1 Hz, 1H), 7.75 (t, *J* = 7.7 Hz, 1H), 7.52 (d, *J* = 4.6 Hz, 1H), 7.44 – 7.37 (m, 1H), 5.00 (s, 2H), 2.70 (s, 3H) ppm. ^13^C NMR (101 MHz, CDCl_3_) δ 197.9, 158.4, 152.0, 150.5, 147.2, 141.3, 137.4, 132.8, 132.0, 129.4, 127.6, 127.1, 124.1, 120.9, 119.8, 40.5, 24.6 ppm.

### 6-methoxy-4-(2-(6-methylpyridin-2-yl)-5,6-dihydro-4H-pyrrolo[1,2-b]pyrazol-3-yl)quinoline (5a)

2-(6-Methoxyquinolin-4-yl)-1-(6-methylpyridin-2-yl)ethan-1-one (**4a**, 3.37 g, 11.5 mmol) and 1-aminopyrrolidin-2-one tosylate salt (3.77 g, 13.8 mmol) were dissolved in a mixture of toluene (44 mL), DMF (10 mL) and 2,6-lutidine (3.4 mL). The resulting solution was heated up in a Dean-Stark for 16 h. Then cesium carbonate (7.52 g, 23.1 mmol) was added and the suspension was stirred 16 h under Dean-Stark conditions. Water (150 mL) was added to the reaction and the aqueous layer was extracted with EtOAc (3×150 mL), dried over anh MgSO_4_ and concentrated *in vacuo*. The crude was purified by column chromatography (eluted with a gradient of hexane and EtOAc from 50:50 to 0:100) to yield 3.29 g (80%) of **5a**. The spectroscopic properties were identical to the ones described^47, 48^; ^1^H NMR (400 MHz, CDCl_3_) δ 8.74 (d, *J* = 4.4 Hz, 1H), 7.98 (d, *J* = 9.2 Hz, 1H), 7.35 – 7.20 (m, 4H), 6.96 – 6.85 (m, 3H), 4.37 (t, *J* = 7.2 Hz, 2H), 3.54 (s, 3H), 2.91 (br s, 2H), 2.75 – 2.65 (m, 2H), 2.38 (s, 3H) ppm. ^13^C NMR (101 MHz, CDCl_3_) δ 158.6, 157.7, 153.7, 151.8, 147.7, 146.7, 144.9, 139.9, 136.4, 131.2, 128.3, 122.4, 122.2, 121.9, 119.5, 110.5, 104.0, 55.5, 48.5, 26.2, 24.6, 23.3 ppm. HRMS (ESI): m/z [M + H]^+^ calculated for C_22_H_20_N_4_O: 356.1637; found: 356.1639.

### 6-bromo-4-(2-(6-methylpyridin-2-yl)-5,6-dihydro-4H-pyrrolo[1,2-b]pyrazol-3-yl)quinoline (5b)

To a stirred solution of 2-(6-bromoquinolin-4-yl)-1-(6-methylpyridin-2-yl)ethan-1-one (**4b**, 19.8 g, 58.0 mmol) in DMF (60 mL), toluene (268 mL) and 2,6-lutidine (20 mL) at rt was added 1-aminopyrrolidin-2-one tosylate salt (18.9 g, 69.6 mmol). The reaction was heated to reflux under Dean-Stark conditions until most of the starting material had been consumed, as indicated by TLC. The mixture was cooled to rt, cesium carbonate (37.8 g, 116 mmol) was added and the mixture heated to reflux. The reaction was monitored by TLC until all reaction intermediate was consumed. Then, toluene was distilled until the reaction mixture reached 145 °C and then cooled to rt. Water was added (630 mL) and the mixture was stirred at 0 °C for 2h. The precipitate was filtered and washed with more water to obtain a brown solid that was purified twice through a silica column (eluent: EtOAc/MeOH from 0 to 5 %) to obtain **5b** as a pale-brown solid (16.1 g, 68%). The spectroscopic properties were identical to the ones described^16^; ^1^H NMR (400 MHz, CDCl_3_) δ 8.85 (d, *J* = 4.4 Hz, 1H), 7.98 (d, *J* = 8.9 Hz, 1H), 7.92 (d, *J* = 2.2 Hz, 1H), 7.71 (dd, *J* = 8.9, 2.2 Hz, 1H), 7.35 (t, *J* = 7.7 Hz, 1H), 7.31 (d, *J* = 4.4 Hz, 1H), 7.11 (d, *J* = 7.8 Hz, 1H), 6.92 (d, *J* = 7.6 Hz, 1H), 4.37 (t, *J* = 7.2 Hz, 2H), 2.90 – 2.84 (m, 2H), 2.75 – 2.66 (m, 2H), 2.24 (s, 3H) ppm. ^13^C NMR (101 MHz, CDCl_3_) δ 158.2, 153.6, 153.2, 151.3, 150.2, 147.1, 146.7, 140.9, 136.3, 132.5, 131.3, 128.7, 128.7, 123.0, 121.8, 120.2, 118.9, 48.3, 26.0, 24.2, 23.2 ppm.

### 4-(2-(6-methylpyridin-2-yl)-5,6-dihydro-4H-pyrrolo[1,2-b]pyrazol-3-yl)quinolin-6-ol (HYL001)

#### Route a. From 5a

A suspension of NaH (60% dispersion in oil, 6.41 g, 160 mmol) in 60 mL of anh. DMF was prepared under N_2_ atm. and 38.4 mL (160 mmol) of dodecanethiol were added to form a thick foam. Then, a solution of 11.4 mg (32.1 mmol) of 6-methoxy-4-(2-(6-methylpyridin-2-yl)-5,6-dihydro-4H-pyrrolo[1,2-b]pyrazol-3-yl)quinoline (**5a**) in 90 mL of anh DMF was added via cannula and the mixture was heated up to 150 °C and stirred for 30 minutes. Then the reaction was cooled to rt, diluted with EtOAc (100 mL) and extracted twice with NaOH (1M, 100 mL). The aqueous extracts were neutralized with HCl and the resulting solid was filtered. The cake was dissolved in 3M HCl and washed with hexanes (40 mL). The aqueous phase was then neutralized with NaOH and the solid filtered and dried to yield 10.3 g (89%) of **1a** (HYL001) as an off-white solid.

#### Route b, from 5b

To a flame-dried flask, 6-bromo-4-(2-(6-methylpyridin-2-yl)-5,6-dihydro-4H-pyrrolo[1,2-b]pyrazol-3-yl)quinoline (**5b**, purified by chromatography, 7.15 g, 17.6 mmol), Pd(dppf)Cl_2_ (644 mg, 0.88 mmol), KOAc (5.19 g, 52.9 mmol) and bis(pinacolato)diboron (5.38 g, 21.2 mmol) were added under N_2_. Degassed dioxane (140 mL) was added and the mixture heated to 90 °C for 3h. The reaction was slowly cooled to 0 °C and diluted with NaOH (5M, 35.3 mL, 176 mmol). Then, a mixture of 30% H_2_O_2_ (18 mL, 176 mmol) and water (36 mL) was added dropwise. After 18h at rt, the reaction was cooled to 0 °C and quenched with sodium sulphite (2M, 83 mL, 176 mmol). After stirring for 30 minutes, the reaction mixture was checked for the presence of peroxides, filtered and concentrated under vacuum. The resulting aqueous solution was neutralized with HCl and cooled down to 0 °C. The solid was filtered, washed with water and dissolved again in NaOH 5M (70 mL). This solution was refluxed with 2 g of activated carbon for 1h, filtered and neutralized with HCl. The resulting solid was filtered, washed with water and dissolved in HCl 1M (35 mL), then treated with a 20% solution of N-acetylcysteine for 5h. The solid was then precipitated with acid, filtered, dissolved in hot MeOH (300 mL) and refluxed with 1 g of activated charcoal for 1h. The solution was filtered hot and the solvent was removed by distillation to the minimum volume. The resulting white suspension was cooled at 0 °C for 2h and filtered to yield 4.81 g (80%) of **1a** (HYL001) as a white solid.

#### Route c. From 5b

6-Bromo-4-(2-(6-methylpyridin-2-yl)-5,6-dihydro-4H-pyrrolo[1,2-b]pyrazol-3-yl)quinoline (**5b**, 9.23 g, 22.8 mmol), Pd_2_(dba)_3_ (522 mg, 0.57 mmol), KOH (3.0g, 45.6 mmol) and di-tert-butyl(2’,4’,6’-triisopropyl-3,4,5,6-tetramethyl-[1,1’-biphenyl]-2-yl)phosphine (tetramethyl di-tBuXPhos) (548 mg, 1.14 mmol) were introduced in a flame-dried Schlenk flask and dissolved in degassed dioxane (19 mL) and degassed water (9.5 mL). The mixture was vigorously stirred at 100 °C for 4.5h, then it was cooled to rt and 40 mL of a 20% NaOH solution were added. After 30 minutes of stirring, the suspension was filtered, the solid was washed with NaOH (20%, 20 mL) and the resulting cake was dissolved in HCl (37%, 5 mL). To this solution, 2 g of activated charcoal were added and the resulting black suspension was stirred for 1h at reflux, cooled down to rt, filtered and washed with water. The aqueous solution was neutralized to pH 7 and filtered. The resulting solid was again dissolved in hot MeOH (450 mL), activated charcoal was added and the suspension was refluxed for 1h. The mixture was filtered hot and the filtrate was distilled to the minimum volume and cooled down to 0 °C for 1h. The off-white solid was filtered, washed with cold MeOH and dried under vacuum to yield 5.45 g (70%) of **1a** (HYL001) as a white solid. The spectroscopic properties were identical to the ones described^47,48^; ^1^H NMR (400 MHz, DMSO) δ 9.66 (s, 1H), 8.58 (d, *J* = 4.4 Hz, 1H), 7.86 (d, *J* = 9.0 Hz, 1H), 7.56 (t, *J* = 7.7 Hz, 1H), 7.48 (d, *J* = 7.8 Hz, 1H), 7.29 – 7.16 (m, 2H), 6.96 (d, *J* = 7.5 Hz, 1H), 6.91 (d, *J* = 2.7 Hz, 1H), 4.27 (t, *J* = 7.2 Hz, 2H), 2.79 (t, *J* = 6.5 Hz, 2H), 2.69 – 2.55 (m, 2H), 1.89 (s, 3H) ppm. ^13^C NMR (101 MHz, DMSO) δ 156.5, 155.1, 152.0, 151.7, 146.6, 146.2, 143.2, 139.4, 136.5, 130.6, 128.7, 122.5, 121.2, 117.7, 109.9, 107.0, 47.9, 25.5, 23.4, 22.5 ppm. IR (film): 3412, 2949, 2833, 1650, 1618, 1508, 1236, 1010 cm^-1^. HRMS (ESI): m/z [M + H]^+^ calculated for C_21_H_18_N_4_O: 342.1481; found: 342.1480.

### Kinase affinity and selectivity

The affinity of the ALK5 inhibitor HYL001 was determined for the ALK kinase family using the KINOMEscan^TM^ Technology—measuring the ability of compounds to competitively inhibit binding between DNA-tagged kinases and immobilized assay ligands, using quantitative PCR—run by Eurofins/DiscoveryX (Birmingham AL, USA). Dissociation constants were calculated by measuring the amount of kinase captured on the solid support as a function of the test compound concentration. K_d_ values were obtained by an 11-point half-log dilution range of 0.5–30,000 nM of inhibitor for ALK1 (ACVRL1), ALK2 (ACVR1), ALK3 (BMPR1A), ALK4 (ACVR1B), ALK6 (BMPR1B), ACVR2B and TGFβR2. In addition, this technology was used to assess selectivity towards a panel of 97 Kinases (scanEDGE) by Eurofins Panlabs (Grapevine TX, USA). The strength of competitive binding, defined as the percentage of binding inhibition of kinases to assay ligand, was visualized using the TREEspot software (DiscoverRx Corporation) and a selectivity score was calculated (the fraction of non-mutated kinases tested with competitive binding inhibition above threshold percentages 65 or 90).

### Enzymatic activity

The activity, measured as IC_50_, of inhibitors was determined by radiometric γ-33P-ATP-mediated in vitro phosphorylation assays, using the Kinase Profiler service from Eurofins Pharma Discovery Services (Wolverhampton, UK) and Eurofins Cerep (Celle L’Evescault, France). Compounds, prepared in 100% DMSO, were tested using a 9-point curve with half-log serial dilutions (1–10,000 nM). For recombinant TGFBR1, the fragment 200–end (T204D) was used; p38alpha was used as a full-length recombinant protein.

### In vitro cellular TGFβ reporter activity assays

HEK293T cells were purchased from the ATCC and cultured in DMEM supplemented with L-glutamine and 10% fetal bovine serum (Life Technologies) at 37°C and 5% CO_2._ The cells were seeded in 24-well plates and transfected with plasmids encoding 12xCAGA-Firefly_Luc and Tk-Renilla_Luc (75 and 10 ng per well, respectively), using polyethylenimine (Polysciences) as transfection reagent. After 7h, medium was replaced for starvation medium (DMEM + 0.05% FBS). The next day, cells were treated with galunisertib, HYL001 or vactosertib, from a 10 mM stock solution in DMSO, in half-log dilutions (0.1–10,000 nM), as well as 5 ng/ml recombinant human TGFB1 (Peprotech). Luciferase activity was measured 16h later using the Dual Luciferase Assay kit (Promega): media were aspirated, and cells were lysed in 200 µl passive lysis buffer (kit) for 20 minutes. Bioluminescence was measured in a Berthold Lumat LB6507 luminometer (18 µl reagents, 10 seconds measurements). Results were normalized to the Renilla transfection control. Inhibition–concentration curves were analysed in Prism Graphpad Prism (v10.1.1) with a least squares regression non-linear fit.

### Plasma Binding Protein

Rapid equilibrium dialysis was performed in triplicates with a rapid equilibrium dialysis (RED) by Sai Life Sciences Lt. (Telangana, India). The RED device contains a dialysis membrane with a molecular weight cut-off of 8,000 Da, separating two chambers for plasma and buffer, respectively. 200 uL of 5 µM warfarin or HYL001 were added to the plasma chamber, versus phosphate buffer saline (pH 7.4) to the buffer chamber. Dialysis was performed for 4h with 100 RPM shaking at 37°C. To assess recovery and stability, aliquots of warfarin or test compounds in plasma were either frozen immediately (T0 sample), or incubated at 37°C for 4h without dialysis (non-dialysed), respectively.

Following dialysis, an aliquot was removed from either chamber and diluted with equal volume of opposite matrix to nullify the matrix effect. Similarly, buffer was added to recovery and stability samples. Next, acetonitrile (containing internal standard, glipizide) was added to the mixtures for protein precipitation and vortexed for 5 minutes. The samples were centrifuged at 4000 RPM at 4°C for 10 min and the supernatant was submitted for liquid chromatography mass spectrometry (LC-MS/MS) analysis. Samples were monitored for parent compound using multiple reaction monitoring (MRM) mode.

The peak area ratios (PAR, analyte versus internal standard) in plasma *vs* bufffer were used to determine the fraction of compound bound to plasma proteins. The following equation was used to determine the extent of plasma protein binding: percent free drug = 100% x <u>(PAR_buffer_</u> <u>/ PAR_plasma_);</u> percent bound drug = 100 - % free drug. Recovery was calculated as 100% x (PAR_plasma_ + PAR_buffer_)_dialysed_ / (PAR_plasma_)_non-dialysed_; stability as 100% x (PAR)_non-dialysed_ / (PAR_plasma_)_frozen at T0_.

### Metabolic stability in human primary hepatocytes

This assay was performed at Sai Life Sciences Lt. (Telangana, India) with human suspension hepatocytes (HMCS1S, Gibco USA). 200 µL of cell suspension containing 2 x 10^6^ hepatocytes/mL (>95% viability) was added to individual wells of 24 well plates and incubated in a CO_2_ incubator at 37°C and 5% CO_2_ for 15 min. T0 incubation was terminated by adding 1000 µL of ice-cold acetonitrile. All other reactions were initiated by adding 200 µL of test compound diluted in Krebs Henseleit Buffer (pH 7.4; pre-warmed at 37°C in CO_2_ incubator) and further incubated for 0, 15, and 60 min before being terminated as the T0 sample. Samples were sonicated for 5 min before centrifuging at 2147 g for 15 min at 4°C, and the supernatant was submitted for analysis by LC-MS/MS, using MRM mode. Testosterone and 7-OH coumarin were used as positive controls related to phase I and phase II metabolism, respectively.

The PARs were used for calculation of metabolic stability. The 0 min area ratio was considered as 100%. The slope of the initial linear range of logarithmic curve of percent remaining versus time (elimination constant K_e_) was used for calculation of half-life: T_1/2_ = ln 2 / K_e_., as well as intrinsic clearance (μL/min/million cells): incubation volume / number of cells (in millions) x K_e_. To calculate into full body context, the values of 120 million cells/g liver and 20 g liver/kg body weight were used.

### *In vitro* safety pharmacology

HYL001 was tested at 10 μM for inhibition of binding and in enzymatic activity of relevant proteins linked to adverse drug reaction risks, using the SafetyScreen 44 service from Eurofins (Celle L’Evescault, France). In each experiment, the respective reference compound was tested concurrently with HYL001, and the data were compared with historical values determined at Eurofins. Binding was defined as the % inhibition of the signal of a radioactively labeled ligand specific for each target, and enzymatic activity as the % inhibition of control enzyme activity. Values are shown as the mean of technical duplicates, error bars represent SD. Results showing an inhibition or stimulation higher than 50% are considered to represent significant effects of the test compound.

The potential degree of hERG inhibition by HYL001 at 3 concentrations was measured by automated patch clamp (Eurofins, Celle L’Evescault, France) on CHO-K1 cells. The tail current amplitude, induced by a one second test pulse to −40 mV after a two second pulse to +20 mV, was measured before and after drug incubation and expressed as % inhibition.

### Mutagenic potential

The Ames test was performed at Xenomatrix (Allschwil, Switzerland). Bacteria from the *S. typhimurium* strains TA98, TA100, TA1535 and TA1537 were exposed to 6 concentrations of a HYL001 (1–320 μg/ml; 3–934 μM; performed in triplicates) in the absence or presence of liver S9 extracts, as well as a positive and a negative control, for 90 minutes in medium containing sufficient histidine to support approximately two cell divisions. After exposure, the cultures were diluted in pH indicator medium lacking histidine or tryptophan and aliquoted into 48 wells of a 384-well plate. Within two days, cells that have undergone reversion to amino acid prototrophy grow into colonies. The number of wells containing revertant colonies were counted by pH indicator colour change for each dose and compared to a solvent (negative) control. A dose dependent increase in the number of revertant colonies upon exposure to HYL001 would have indicated mutagenicity.

### Animal Welfare

All procedures involving animal experimentation conducted at Sai Life Sciences Lt. were carried out in accordance with the guidelines provided by the Committee for the Purpose of Control and Supervision of Experiments on Animals (CPCSEA), and after approval by the Institutional Animal Ethics Committee (IAEC). The study procedures and husbandry care of the study animals were performed in compliance with Association for Assessment and Accreditation of Laboratory Animal Care (AAALAC) (Unit No. 001384) and CPCSEA (Reg. No. 2121/PO/Rc/S/21/CPCSEA) norms.

All experiments involving animal experimentation conducted at IRB Barcelona were approved by the Animal Care and Use Committee of Barcelona Science Park and the Catalan Government (protocol 9162). Mice were maintained in a specific-pathogen-free (SPF) facility with a 12-h light–dark cycle, under controlled temperature and humidity (18-23°C and 40-60% respectively) and given ad libitum access to standard diet and water. All mice were closely monitored by authors, facility technicians (during treatments) and by an external veterinary scientist responsible for animal welfare.

### Pharmacokinetics of HYL001 in BALB/c Mice and Sprague Dawley Rats

The plasma pharmacokinetics of HYL001 in male BALB/c mice and male Sprague Dawley rats were investigated following a single oral administration at 71 mg/kg dose by Sai Life Sciences Lt. (Telangana, India). A total of nine mice were used in this study with 3 mice/time point, following a sparse sampling design. Similarly, three rats were used in this study. Animals were administered orally with a suspension formulation of HYL001 at 71 mg/kg dose. The formulation vehicle was 1% *v/v* NaCMC, 0.4% *v/v* SLS and 0.085% *v/v* PVP 40T in RO water.

Blood samples (approximately 60 μL for mice and approximately 120 μL for rats) were collected under light isoflurane anesthesia (Surgivet) from retro orbital plexus from a set of three mice at Pre-dose, 0.25, 0.5, 1, 2, 4, 6, 8, and 24 hr. Immediately after blood collection, plasma was harvested by centrifugation at 4000 rpm, 10 min at 4^0^C and samples were stored at −70 ±10°C until bioanalysis.

All samples were processed for analysis by protein precipitation method and analyzed by LC-MS/MS. The plasma pharmacokinetic parameters were estimated using non-compartmental analysis tool of Phoenix® WinNonlin software (Version 8.0)

### Toxicology Studies in Rats and Mice

Sprague-Dawley rats were procured from Hylasco Bio-Technology Pvt. Ltd., Hyderabad. After completion of the acclimatization period twelve healthy rats were randomly allocated to the control and treatment groups.

Treatment groups included controls, and either HYL001 (25 mg/kg BID or 77 mg/kg/BID) or Vactosertib (82 mg/kg BID) treated animals, having three rats/sex/group. Rats from control group received vehicle, consisting of 0.4 % Sodium Lauryl Sulphate (SLS), 0.085 % Povidone (Polyvinylpyrrolidone), 0.05% Antifoam A and quantity sufficient to 1% w/v Carboxy Methyl Cellulose Sodium Salt. Rats from treatment groups were administered with suspensions of the indicated drugs in vehicle, orally twice a day for an intended period of 28 consecutive days. The dosing volume was kept constant at 10 mL/kg for each rat. Rats that showed any critical clinical sign that worsened during the treatment, were necropsied in the shortest time for ethical reasons. The rest of the rats were necropsied at the end of the treatment.

Following 28 days of treatment, or when found sick, the rats were humanely euthanized by carbon dioxide asphyxiation on day 29, except for one group of HYL001 treated rats with 71 mg/kg/BID, that were allow to recover from treatment for further 28 days.

The rats were subjected to detailed gross pathological examination which included careful examination of the external surface of the body, all orifices and the cranial, thoracic and abdominal cavities and their contents. After gross pathological examination the vital organs were trimmed of any adherent tissue and were weighed wet. Paired organs were weighed together. Organ weights relative to terminal body weights were calculated for each rat.

Parameters evaluated during the study included in-life observations such as clinical signs observation, body weights, percent body weight gains, and feed consumption. Parameters like hematology, clinical chemistry, gross pathology, and organ weights were evaluated on the day of termination. Several organs/tissues were further processed and evaluated for histopathology, where special attention was given to bones, gastrointestinal system, heart, and vascular lesions in the aorta and coronary arteries. For histopathology, 3-5 μM thick tissue sections were cut and stained with hematoxylin-eosin stain. The heart was trimmed by longitudinally bisecting along a plane perpendicular to the plane of the pulmonary artery to expose the right atrioventricular, left atrioventricular, and aortic valves. Both halves of the heart were embedded as hemisection in paraffin with the cut surface down. Blocks were sectioned to obtain at least three heart valves.

All the individual animal data were summarized in terms of group mean and standard deviation. Body weight, body weight gain, hematology, clinical chemistry, organ weight data of toxicity group rats were analyzed using “t” test. All analysis and comparisons were evaluated at 5% level i.e. P ≤ 0.05. The statistical analysis was performed using GraphPad Prism statistical software version 5.02 for Windows, GraphPad Software, San Diego California USA.

The study design in mice was similar as in rats, except 5 mice of each sex were used (n=10 in total), and that there was not a recovery group. All analyses were performed at day 29 after termination of treatment.

### Mouse Injections

For all tumour cell injections, C57BL/6J (or athymic BALB/C *nu/nu*) mice were purchased from Janvier at 6 weeks of age and injected at 7–8 weeks. Sex was matched with the origin of the tumour organoids, i.e. males for this study. Intrasplenic or portal vein injections, respectively with 31G needles on 1 ml insulin syringes or 33G needles on a 50/100 μL Hamilton syringe, were used for liver colonization by the introduction of dissociated organoids (single cells) into the portal circulation. MTOs were cultured in standard six-well plates for four days and trypsinized as described before^5^. The resulting single cell suspension in HBSS (Lonza) was filtered through 100 and 40 μm meshes (to remove clumps of cells and aggregated debris). Intrasplenic injections were performed as previously described^5,8,9^, using 2 x 10^5^ cells in 70 μL HBSS. For portal vein injections, 2.5 x 10^3^ cells in 25 μL HBSS were injected directly into the portal vein^5^. Mice were euthanized at 3–5 weeks or at humane endpoint (advanced metastasis-associated morbidity causing or threatening to cause severe suffering) to obtain survival-type data. Visible liver metastases were counted and data were analysed using GraphPad Prism software (v.7.03).

### Mouse Treatment

HYL001, galunisertib, vactosertib or control vehicle were administered by gavage twice per day, starting at the indicated timepoints after cell injection. Gavage treatments were performed by technicians from the animal facility, not further involved in this study (blinded). For checkpoint immunotherapy or dual treatments, we used rat anti-PD-1 (RMP1-14; Leinco P372) or rat IgG2a (Leinco I-1177) isotype-control antibodies.

### In vivo study design

Experimental group sizes were practically associated to cage sizes (5 mice/cage) and experiments were designed to have n >= 5 per group (1 or more cages) and repeated at least once in independent experiments as in^5^. No mice were excluded from the analysis. For gavage treatment, as control vehicle and compound containing vials were visually distinguishable, the only randomization we performed was the order of injecting mice: researcher performing the injections was blinded to treatment group. End point criteria are equivalent to those described above.

## AUTHOR INFORMATION

### Author Contributions

DT and DB conceived the study; DT, EB, AR, and ES coordinated experiments and wrote the manuscript. AR, DB, JM and CSZ designed, synthesized and analyzed all chemicals, including HYL001. DT determined cellular enzyme activity *in vitro*. IR coordinated all analyses performed by CROs, and together with DT and ES extracted results from reports. NP and IR evaluated toxicity in mice. DT, MC, SP, AH, XHM designed and performed *in vivo* efficacy experiments. MS performed IHC. DT and ES performed visualization of data.

### Notes

DT, DB, JM, AR, and EB hold a patent on the synthesis and use of HYL001^48^. E.B. is author in a patent describing bispecific antibodies to target cancer stem cells. The laboratory of E.B. has received research funding from MERUS and INCYTE. E.B. has received honoraria for consulting from Genentech.

## ACKNOWLEDGEMENTS

This work was supported by grants to EB from the Spanish Ministry of Science / European Next generation Funds (PDC2021-121226-I00); the BioMedTec programme from “la Caixa” Foundation (ID 100010434), under the agreement LCF/PR/GN18/50310010, and two Gínjol grants (2020-07-012 and 2021-08-011) from the Catalan Government. This work was also supported by grants to AR from FEDER/Ministerio de Ciencia e Innovación (MICINN) (PID2020-115074GB-I00/AEI/10.13039/501100011033) and Generalitat de Catalunya (2021 SGR 00866). IRB Barcelona is the recipient of institutional funding from MICINN through the Centres of Excellence Severo Ochoa Award and from the CERCA Program of the Catalan Government.

## REFERENCES

1. Sung, H. et al. Global Cancer Statistics 2020: GLOBOCAN Estimates of Incidence and Mortality Worldwide for 36 Cancers in 185 Countries. CA Cancer J Clin 71, 209–249 (2021).

2. Galon, J. & Bruni, D. Approaches to treat immune hot, altered and cold tumours with combination immunotherapies. Nat Rev Drug Discov 18, 197–218 (2019).

3. Janssen, E., Subtil, B., de la Jara Ortiz, F., Verheul, H. M. W. & Tauriello, D. V. F. Combinatorial Immunotherapies for Metastatic Colorectal Cancer. Cancers (Basel) 12, 1875 (2020).

4. Batlle, E. & Massagué, J. Transforming Growth Factor-β Signaling in Immunity and Cancer. Immunity 50, 924–940 (2019).

5. Tauriello, D. V. F. et al. TGFβ drives immune evasion in genetically reconstituted colon cancer metastasis. Nature 554, 538–543 (2018).

6. Tauriello, D. V. F., Sancho, E. & Batlle, E. Overcoming TGFβ-mediated immune evasion in cancer. Nat Rev Cancer 22, 25–44 (2022).

7. Derynck, R., Turley, S. J. & Akhurst, R. J. TGFβ biology in cancer progression and immunotherapy. Nat Rev Clin Oncol 18, 9–34 (2021).

8. Calon, A. et al. Dependency of Colorectal Cancer on a TGF-β-Driven Program in Stromal Cells for Metastasis Initiation. Cancer Cell 22, 571–584 (2012).

9. Calon, A. et al. Stromal gene expression defines poor-prognosis subtypes in colorectal cancer. Nat Genet 47, 320–329 (2015).

10. Mariathasan, S. et al. TGFβ attenuates tumour response to PD-L1 blockade by contributing to exclusion of T cells. Nature 554, 544–548 (2018).

11. Peng, D., Fu, M., Wang, M., Wei, Y. & Wei, X. Targeting TGF-β signal transduction for fibrosis and cancer therapy. Molecular Cancer 2022 21:1 21, 1–20 (2022).

12. Kim, K. K., Sheppard, D. & Chapman, H. A. TGF-β1 Signaling and Tissue Fibrosis. Cold Spring Harb Perspect Biol 10, (2018).

13. Kim, B.-G., Malek, E., Choi, S. H., Ignatz-Hoover, J. J. & Driscoll, J. J. Novel therapies emerging in oncology to target the TGF-β pathway. J Hematol Oncol 14, 55 (2021).

14. Anderton, M. J. et al. Induction of heart valve lesions by small-molecule ALK5 inhibitors. Toxicol Pathol 39, 916–924 (2011).

15. Yingling, J. M. et al. Preclinical assessment of galunisertib (LY2157299 monohydrate), a first-in-class transforming growth factor-β receptor type I inhibitor. Oncotarget 9, 6659–6677 (2018).

16. Li, H.-Y. et al. Optimization of a dihydropyrrolopyrazole series of transforming growth factor-beta type I receptor kinase domain inhibitors: discovery of an orally bioavailable transforming growth factor-beta receptor type I inhibitor as antitumor agent. J Med Chem 51, 2302–6 (2008).

17. Herbertz, S. et al. Clinical development of galunisertib (LY2157299 monohydrate), a small molecule inhibitor of transforming growth factor-beta signaling pathway. Drug Des Devel Ther 9, 4479–99 (2015).

18. Jin, C. H. et al. Discovery of N-((4-([1,2,4]triazolo[1,5-a]pyridin-6-yl)-5-(6-methylpyridin-2-yl)-1H-imidazol-2-yl)methyl)-2-fluoroaniline (EW-7197): a highly potent, selective, and orally bioavailable inhibitor of TGF-β type I receptor kinase as cancer immunotherapeutic/antifibrotic agent. J Med Chem 57, 4213–38 (2014).

19. Mathijssen, R. H. et al. Clinical pharmacokinetics and metabolism of irinotecan (CPT-11). Clin Cancer Res 7, 2182–94 (2001).

20. Mancarella, S. et al. Validation of hepatocellular carcinoma experimental models for TGF-β promoting tumor progression. Cancers (Basel) 11, (2019).

21. Naka, K. et al. Novel oral transforming growth factor-β signaling inhibitor EW-7197 eradicates CML-initiating cells. Cancer Sci 107, 140–148 (2016).

22. Bowes, J. et al. Reducing safety-related drug attrition: The use of in vitro pharmacological profiling. Nat Rev Drug Discov 11, 909–922 (2012).

23. Mathes, C. QPatch: The past, present and future of automated patch clamp. Expert Opin Ther Targets 10, 319–327 (2006).

24. Stauber, A. J., Credille, K. M., Truex, L. L., Ehlhardt, W. J. & Young, J. K. Nonclinical Safety Evaluation of a Transforming Growth Factor β Receptor I Kinase Inhibitor in Fischer 344 Rats and Beagle Dogs. J Clin Toxicol 4, 1–10 (2014).

25. Frazier, K. et al. Inhibition of ALK5 signaling induces physeal dysplasia in rats. Toxicol Pathol 35, 284–295 (2007).

26. Martin, C. J. et al. Selective inhibition of TGFβ1 activation overcomes primary resistance to checkpoint blockade therapy by altering tumor immune landscape. Sci Transl Med 12, eaay8456 (2020).

27. Cañellas-Socias, A. et al. Metastatic recurrence in colorectal cancer arises from residual EMP1+ cells. Nature 611, 603–613 (2022).

28. Álvarez-varela, A. et al. Mex3a marks drug-tolerant persister colorectal cancer cells that mediate relapse after chemotherapy. 3, (2022).

29. Nakanishi, Y. et al. Simultaneous Loss of Both Atypical Protein Kinase C Genes in the Intestinal Epithelium Drives Serrated Intestinal Cancer by Impairing Immunosurveillance. Immunity 49, 1132–1147.e7 (2018).

30. Kasashima, H. et al. Stromal SOX2 Upregulation Promotes Tumorigenesis through the Generation of a SFRP1/2-Expressing Cancer-Associated Fibroblast Population. Dev Cell 56, 95–110.e10 (2021).

31. Peuker, K. et al. Microbiota-dependent activation of the myeloid calcineurin-NFAT pathway inhibits B7H3-and B7H4-dependent anti-tumor immunity in colorectal cancer. Immunity 55, 701–717.e7 (2022).

32. Martinez-Ordoñez, A. et al. Hyaluronan driven by epithelial aPKC deficiency remodels the microenvironment and creates a vulnerability in mesenchymal colorectal cancer. Cancer Cell 41, 252–271.e9 (2023).

33. Jung, G., Benítez-Ribas, D., Sánchez, A. & Balaguer, F. Current Treatments of Metastatic Colorectal Cancer with Immune Checkpoint Inhibitors-2020 Update. J Clin Med 9, 1–14 (2020).

34. Tschernia, N. P. & Gulley, J. L. Tumor in the Crossfire: Inhibiting TGF-β to Enhance Cancer Immunotherapy. Biodrugs 36, 153 (2022).

35. Di, L. An update on the importance of plasma protein binding in drug discovery and development. Expert Opin Drug Discov 16, 1453–1465 (2021).

36. Gerritse, S. L. et al. High-dose administration of tyrosine kinase inhibitors to improve clinical benefit: A systematic review. Cancer Treat Rev 97, (2021).

37. Jung, M. et al. 1453P Safety and efficacy of vactosertib, a TGF-βR1 kinase inhibitor, in combination with paclitaxel in patients with metastatic gastric adenocarcinoma. Annals of Oncology 31, S912 (2020).

38. Malek, E. et al. Vactosertib, a novel TGF-β1 type I receptor kinase inhibitor, improves T-cell fitness: a single-arm, phase 1b trial in relapsed/refractory multiple myeloma. Res Sq 142, 4749–4749 (2023).

39. Ahn, J.-H. et al. Clinical Activity of TGF-β Inhibitor Vactosertib in Combination with Imatinib in Desmoid Tumors: A Multicenter Phase Ib/II Study. Clinical Cancer Research OF1–OF9 (2024) doi:10.1158/1078-0432.CCR-23-2823.

40. Nair, A. B. & Jacob, S. A simple practice guide for dose conversion between animals and human. J Basic Clin Pharm 7, 27 (2016).

41. Le, D. T. et al. Mismatch repair deficiency predicts response of solid tumors to PD-1 blockade. Science (1979) 357, 409–413 (2017).

42. Lind, H. et al. Dual targeting of TGF-β and PD-L1 via a bifunctional anti-PD-L1/TGF-βRII agent: status of preclinical and clinical advances. J Immunother Cancer 8, (2020).

43. Lan, Y. et al. Enhanced preclinical antitumor activity of M7824, a bifunctional fusion protein simultaneously targeting PD-L1 and TGF-β. Sci Transl Med 10, eaan5488 (2018).

44. Mundla, S. R. A pyridin quinolin substituted pyrrolo [1,2-b] pyrazole monohydrate as tgf-beta inhibitor. WO 2007/018818 A1. (2007).

45. Kim, D.-K. et al. 2-pyridyl substituted imidazoles as therapeutic ALK5 and/or ALK4 inhibitors. WO 2012/002680 A2. (2012).

46. Campbell, K. N. & Schaffner, I. J. The Preparation of 4-Methylquinolines. J Am Chem Soc 67, 86–89 (1945).

47. De, K., Legros, J., Crousse, B. & Bonnet-Delpon, D. Solvent-promoted and -controlled aza-Michael reaction with aromatic amines. J Org Chem 74, 6260–6265 (2009).

48. Tauriello, D., Byrom, D., Matarín-Morales, J. A., Batlle-Gómez, E. & Riera-Escalé, A. TGFβ INHIBITOR AND PRODRUGS. WO 2020/104648 A2. (2020).

